# A differentiated and durable allogeneic strategy applicable to cell therapies

**DOI:** 10.1101/2025.03.27.645807

**Authors:** Utsav Jetley, Ishina Balwani, Palak Sharma, Ian Miller, Allie Luther, Ivy Dutta, Nithila Saravanan, Qingzhan Zhang, Boning Zhang, Ozgun Kilic, Biao Liu, Bo W Han, Dai Liu, Birgit Schultes, Aaron Prodeus, Yong Zhang

**Affiliations:** Intellia Therapeutics, Inc, Cambridge, MA, USA

**Keywords:** Allogeneic T cell therapy, CAR-T, TCR-T, CRISPR, iPSC

## Abstract

Autologous T cell therapies have shown profound clinical responses; however, their widespread use has been limited primarily due to their individualized manufacturing requirements. To develop a persistent "off-the-shelf" allogeneic (Allo) approach, a multiplex Nme2Cas9-based cytosine base editor was deployed to knockout select HLA Class I and II alleles (*HLA-A*, *HLA-B*, and the class II transactivator (*CIITA*)), while retaining HLA-C to protect from NK cell rejection. Matching the residual HLA-C allele from homozygous donors to the host prevented rejection of the donor T cells by allogeneic host T and NK cells. Site-specific integration of a tumor-specific CAR or TCR into the *TRAC* locus using SpyCas9 nuclease and an adeno-associated virus (AAV) template allowed for high localized insertion rate while simultaneously removing the endogenous TCR and preventing GvHD. Using a lipid nanoparticle (LNP)-based delivery system of the editing components enabled a robust cell engineering process, achieving high editing rates and cell expansion. These allogeneic T cells demonstrated comparable functional activity to their autologous counterparts in preclinical assays. Moreover, this gene editing approach generated cells with minimal chromosomal aberrations. The Allo strategy has also been applied to induced pluripotent stem cells (iPSCs), suggesting potential applications in regenerative medicine applications.

## Introduction

Patient-derived autologous chimeric antigen receptor (CAR) T cell therapies have shown remarkable success for treating several types of hematological malignancies. Currently, the United States Food and Drug Administration (FDA) has approved five CD19-targeted CAR T and two B cell maturation antigen (BCMA)-targeted CAR T cells. ^1,2^ Besides hematological malignancies, CAR-T therapy also shows encouraging early efficacy in solid tumors. For example, several reports have demonstrated the promising efficacy of GD2-CAR T cell therapy for H3K27M mutated diffuse midline gliomas ^3,4^ and for high-risk neuroblastoma. ^5,6^ A recently completed first-in-human trial showed that repetitive intracerebroventricular dosing of B7-H3 CAR T cells in pediatric and young adult patients with diffuse intrinsic pontine glioma is tolerable, including multiyear repeated dosing, and may have clinical efficacy.^7^ GPC3 CAR-T cells armored with IL-15 or dominant negative TGFβRII showed encouraging anti-tumor activity in patients with hepatocellular carcinoma. ^8,9^ The transformative impact of CAR T cell therapy on B cell cancers has inspired researchers to explore its potential in treating autoimmune diseases. Emerging clinical trials of CD19-targeted or BCMA-targeted CAR T cells have shown promising early results in patients with B cell-driven autoimmune conditions, including systemic lupus erythematosus, ^10,11^ idiopathic inflammatory myopathies, ^12^ immune-mediated necrotizing myopathy, ^13^ myasthenia gravis, ^14–16^ neuromyelitis optica spectrum disorder, ^17^ and systemic sclerosis. ^18^

In addition to CAR-T therapy, other forms of autologous T cell therapies have shown remarkable progress in treating solid tumors, including tumor-infiltrating lymphocytes (TIL) therapy and T cell receptor (TCR) T cell therapy. For example, in August 2024, the FDA granted accelerated approval to a TCR-T cell therapy targeting MAGE-A4 in synovial sarcoma, a rare type of soft tissue cancer. ^19^

Despite the impressive efficacy of T cell therapies, several challenges have hindered their widespread adoption. Firstly, the individualized nature of autologous T cell therapy results in significantly higher cost compared to other modalities, ranging from 373,000 to 475,000 USD per dose in the United States. ^20^ Secondly, oncology patients who have undergone multiple lines of treatment, such as radiotherapy and chemotherapy, often have T cells with poor quality, leading to manufacturing failures and compromised efficacy underscoring the critical importance of high-quality T cells as starting material. Additionally, a case study reported a rare event, where the resistance to CAR T cell therapy occurred due to contaminating tumor cells in the T cell product derived from a leukemia patient. ^21^ Lastly, autologous CAR-T cell production faces complex manufacturing and logistical issues associated with limited manufacturing slots, manufacturing failures, and long production time. As a result, these therapies are less accessible to patients with aggressive disease who succumb to their illness during the manufacturing process. This is consistent with a recent analysis, which showed that, among 41% of community-referred non-Hodgkin lymphoma patients who were unable to access anti- CD19 CAR T-cell therapy, most of them were due to disease progression (34%) or poor health (15%) prior to treatment. ^22^

Given the significant challenges of scale, cost, and complexity associated with autologous T cell therapy, there is a strong desire for an immediately available cell therapy option. However, development of “off-the-shelf” CAR-T cells is challenging due to three critical and complex hurdles related to alloreactive immunological responses. ^23^ The main safety concern for T cell products is graft vs host disease (GvHD) that can be mediated by the donors’ T cell receptors reacting with foreign human leukocyte antigens (HLA) or presented antigens derived from the patient’s (host) healthy tissues. This “non-self” recognition will initiate an attack on the patient’s body by the donor allogeneic T cells, which leads to GvHD, if not addressed.

This issue has been largely mitigated by knocking out TCRs in donor T cells via various gene editing approaches or the use of virus-specific T cells. ^24,25^ Inversely, due to the similar HLA and TCR interaction mechanism, T cells from the recipient patient will recognize allogeneic T cells as “non-self” if they express mismatched HLA alleles and initiate the “defense” machinery to eliminate those foreign T cells. Removing all HLA molecules from the allogeneic donor, for example by CRISPR/Cas9 knockout of beta-2 microglobulin (B2M), to evade T cell-mediated recognition by the recipient is being investigated. ^26^ However, this approach induces a different challenge, as the patient’s natural killer cells (NKs) from the innate immune system will recognize these allogeneic donor T cells with low or no HLA class I expression (“missing self”) and rapidly reject the therapeutic T cells. ^27^ Both host T cell and NK cell mediated immune rejection leads to limited allogeneic T cell persistence, which in turn compromises efficacy, particularly response duration, of these therapeutic T cells.

In this study, we present the development of a differentiated allogeneic T cell platform that harnesses the power of our orthogonal CRISPR/Cas9 cleavase and base editor systems, coupled with lipid nanoparticle (LNP) / adeno-associated virus (AAV) delivery technologies. To expand the applications of our allogeneic platform, we extended our research to iPSC-derived pancreatic progenitors and cardiomyocytes. Our data suggest that this allogeneic platform is protected from T and NK cell rejection and readily deployable for TCR- and CAR-T cell therapy, and potentially iPSC-derived regenerative cell therapies.

## Results

### Overview of the Allo platform

To develop an effective Allo platform, we need to address three key immunological challenges associated with Allo T cell therapies (Fig. 1A). The first is GvHD as Allo T cells can recognize host antigens via their TCRs. To limit the occurrence and severity of GvHD, we used CRISPR/Cas9 to knock out endogenous TCRs and insert a therapeutic CAR or TCR into the *TRAC* locus by site-specific homology directed repair (HDR)-mediated insertion. This approach prevents allogeneic T cells from recognizing host antigens through their native TCRs and confers therapeutic targeting. The next challenge is host vs graft rejection, which involves both host T cells and host NK cells. To mitigate host T cell rejection, we selectively knocked out HLA-A and HLA-B, but retained HLA-C expression, rather than eliminate all HLA class I as is the case with B2M knockout, to minimize rejection by NK cells. To further prevent host CD8+ T cell activation, we matched donor T cell’s residual HLA-C with host’s HLA-C. Importantly, retention of HLA-C and HLA-E can suppress host NK cell activation, with HLA-C being reported as one of the strongest inhibitors of NK cells. ^28^ Previous studies suggest that host CD4+ T cells can be activated through either the semi-direct pathway, in which T cells recognize intact donor HLA class II molecules and bound peptides displayed on recipient APCs, or the indirect pathway, in which CD4 T cells recognize donor peptides processed and presented by HLA class II molecules in recipient APCs. ^29^ Once activated, CD4+ T cells can further promote acute rejection by helping host CD8+ T cell activation or by facilitating B cell production of donor-specific HLA antibodies, leading to antibody-mediated rejection. ^29,30^ Because T cells upregulate HLA class II upon activation, we also knocked out CIITA to disrupt HLA class II expression in Allo T cells and prevent host CD4 T cell activation. By applying this approach, we aimed to minimize both host T cell and NK cell mediated rejection of the allogeneic cell product (Fig. 1B). To address the notorious *HLA-A* and *HLA-B* polymorphism challenge, we leveraged bioinformatic analysis to identify one lead HLA-A targeting sgRNA and one lead HLA-B targeting sgRNA, which cover 84% of *HLA-A* alleles and 74% of *HLA-B* alleles, respectively, including the most common alleles (data not shown).

**Fig. 1.**
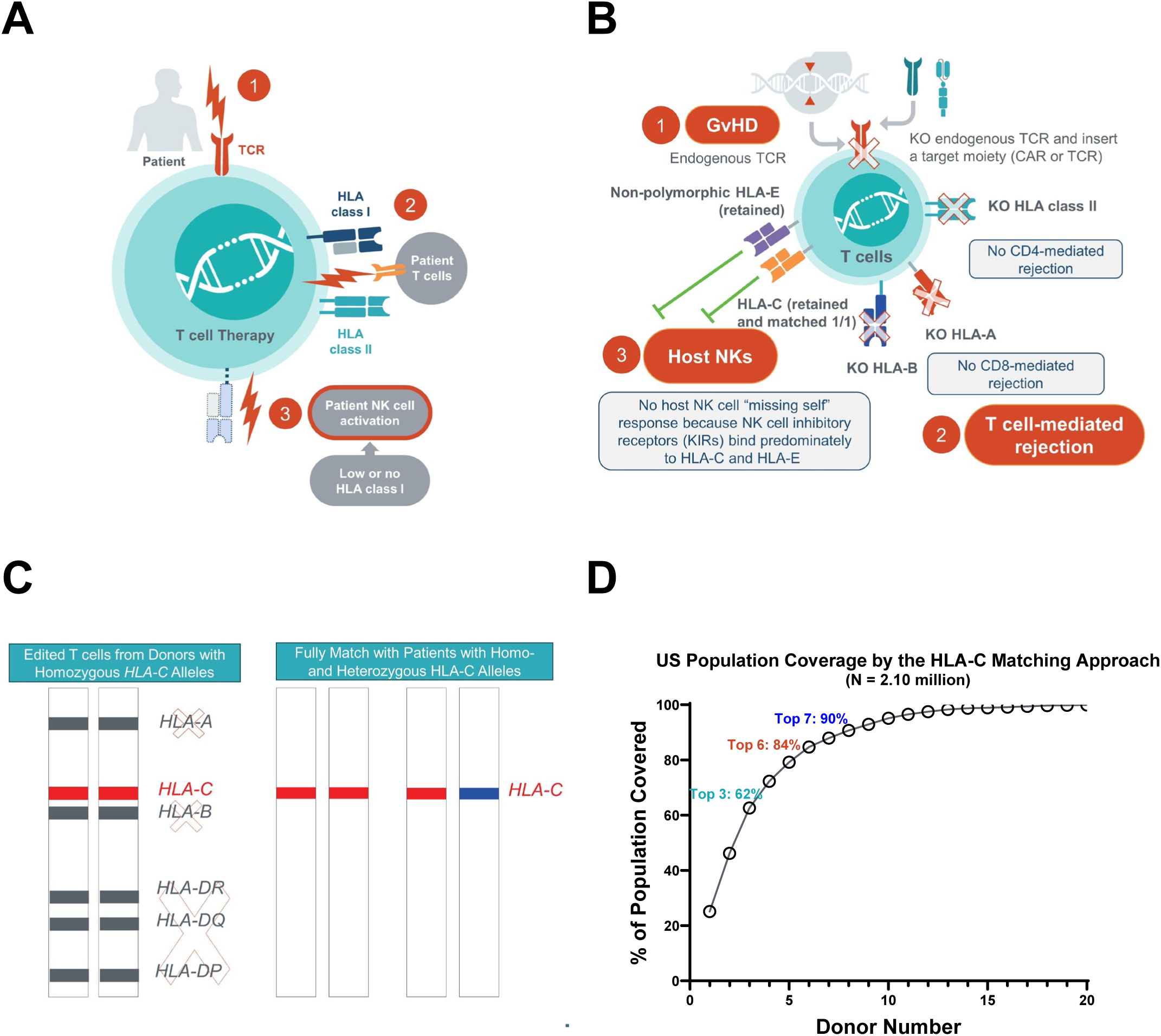
Overview of the Allo platform. A. Schematic of three immune concerns to be addressed by allogeneic T cell therapies. B. Schematic of this differentiated allogeneic approach. C. Schematic of the HLA-C matching strategy which achieves full HLA match by 1 out of 10 HLA allele matching. D. Leveraging a large sample size of over 2 million donors from a US cohort, bioinformatic analysis predicted the population coverage of HLA-C matching using HLA-C homozygous donors.

Traditionally, a 10 out of 10 HLA match (HLA-A, -B, -C, -DR, and -DQ) is considered the gold standard for unrelated donor selection in hematopoietic stem cell transplantation. However, due to the extensive polymorphism of HLA genes, finding a fully matched unrelated donor is very challenging. Because we knock out HLA-A, HLA-B, and HLA Class II, we focus solely on matching HLA-C. To further simplify the matching process, we specifically selected HLA-C homozygous donors so that we can achieve a full match by only matching a single HLA-C allele, effectively transforming the traditional 10/10 match into a simple 1/10 match (Fig. 1C). To understand the feasibility of this 1 out of 10 matching approach, we performed a bioinformatic prediction using one large data set from BeTheMatch containing HLA allelic data from more than 2 million US donors on a per donor basis rather than just allele population frequencies. The large sample size from this cohort gives us strong statistical power to predict top HLA-C genotypes and their coverage in the US population as well as across US ethnicities. Our analysis predicted that 62% of the US population can be covered by the top 3 HLA-C types, while ∼90% coverage can be achieved with the top 7 HLA-C types (Fig. 1D). Leveraging this cohort, we also performed a subgroup analysis to understand the number of HLA-C types needed across different ethnicities, such as Caucasian, African American, Hispanic, Asian etc., which showed comparable coverage across various US ethnicities albeit with slightly different HLA-C rank orders (Sup Fig. 1). Thus, this 1 out of 10 matching can significantly reduce the time and resources required to find suitable donors and increase the likelihood of finding a match for patients from diverse ethnic backgrounds, which is critical to enable a successful Allo platform.

### T Cell Engineering

Efficient and scalable multiplex gene editing with minimal genotoxicity concern is required to enable the described allogeneic platform. CRISPR/Cas9 has been leveraged for high-efficiency gene knockout and insertion in healthy donor T cells. However, traditional approaches for ex-vivo cell engineering including delivery of editing components by electroporation (EP), and simultaneous or repetitive dsDNA breaks by CRISPR/Cas9 nucleases, limit the number of edits feasible of being introduced to T cells without impacting cell viability and introducing genotoxic events such as chromosomal translocations or loss. Additionally, semi-random integration of CAR transgenes from lentiviral vectors adds an additional layer of genotoxicity risk. To mitigate these limitations, we have developed a gene editing platform where the CAR transgene is inserted into the *TRAC* locus using a SpyCas9 cleavase and HDR, while multiple gene knockouts are achieved using a Nme2Cas9 cytosine base editor (CBE) without induction of dsDNA breaks (Sup Fig. 2A). By leveraging two Cas9 orthologs (Nme2Cas9 and SpyCas9) which bind to guide RNAs (gRNAs) with different scaffold sequences, these edit types can occur simultaneously while avoiding cross-editing between intended cleavase/indel and base-editing outcomes (Sup Fig. 2B). Additionally, we found that these gene editing components can be delivered by lipid nanoparticle (LNP) transfection of mRNA and single guide RNA (sgRNA), which can circumvent toxicity and scalability challenges associated with EP. ^31^

**Fig. 2.**
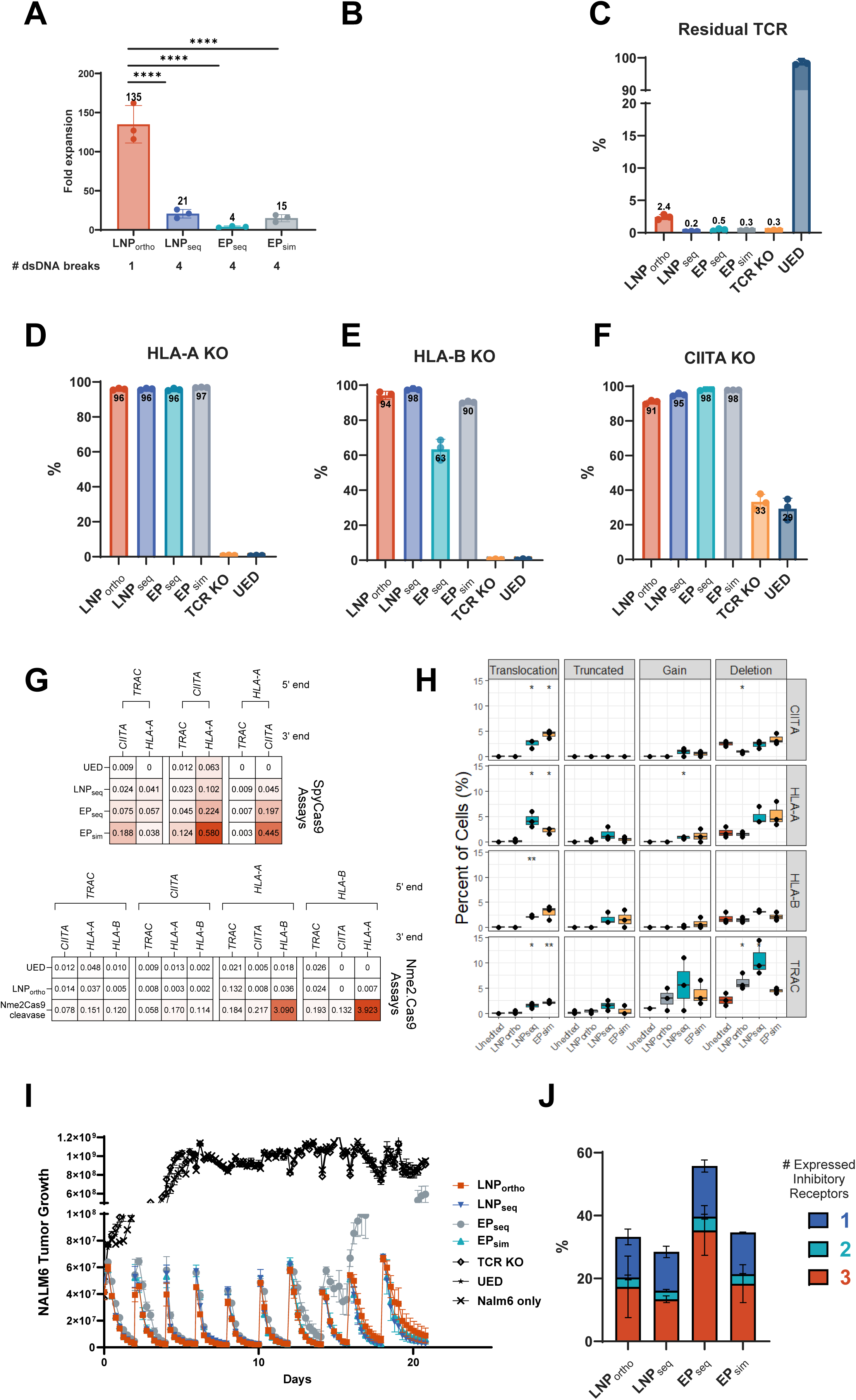
T cell engineering. Allogeneic CD19 CAR T cells were engineered from three healthy donors by either simultaneous electroporation of SpyCas9 RNPs (EP_sim_); sequential electroporation of SpyCas9 RNP (EP_seq_); sequential LNP transfection of SpyCas9 mRNA/sgRNAs (LNP_seq_) or LNP delivery of Nme2Cas9 CBE gene knockouts and CAR insertion using SpyCas9 (LNP_ortho_). A. The fold cell expansion through the engineering process was determined by cell counts and calculating the yield of viable cells on Day 10 vs initial starting material (Day 0). B-F. Editing rates including CAR insertion, TCR knockout (residual TCR expressing cells), HLA-A KO, HLA-B KO, and CIITA KO were quantified by flow cytometry. TCR KO T cells and unedited (UED) T cells served as controls. G. Heatmap of ddPCR assays which were used to quantify the occurrence of translocations across targets, showing a significant reduction of translocations in LNP_ortho_ edited cells with rates at or near background levels of UED cells. H. Chromosomal structural variants were characterized by dGH In-Site Assay™ using unique fluorescent probes bracketing each edit site. Graphs shows levels of translocations, truncations, gain (of probe signal), or large deletion (loss of probe), confirming an overall improved genotoxicity profile with LNP_ortho_ engineered cells vs other methods. I. The allo-CD19 CAR-T cells were evaluated for long-term killing in a serial rechallenge assay with GFP-expressing Nalm6 tumor cells, showing high levels of tumor control in all groups except EP_seq_ edited cells. J. Flow cytometry was used to evaluate expression of exhaustion markers (PD-1, TIM-3, LAG-3) after three rechallenge cycles. 1 stands for only 1 detected exhaustion marker in Allo T cells; 2 stands for 2 detected exhaustion markers in Allo T cells, and 3 stands for 3 detected exhaustion markers in Allo T cells. CBE: cytosine base-editor. \**p*<0.05, \*\**p*<0.01, *** *p*<0.001, \*\*\*\**p*<0.01.

To characterize the potential advantages of this engineering platform, allo-CD19 CAR-T cells with CD19-CAR *TRAC* insertion, HLA-A KO, HLA-B KO, and CIITA KO were engineered from three healthy donors by either 1) EP_sim_: simultaneous electroporation of SpyCas9 RNP, 2) EP_seq_: sequential electroporation of SpyCas9 RNP, 3) LNP_seq_: sequential LNP transfection of SpyCas9 mRNA/sgRNAs, and 4) LNP_ortho_: LNP delivery of Nme2Cas9 CBE gene knockout and CAR insertion using SpyCas9. While viable cells were obtained from all processes, there was a dramatic increase in cell expansion/yield in the LNP_ortho_ group and a particularly poor yield in the EP_seq_ group (Fig. 2A). Despite the higher expansion, flow cytometry showed that CAR insertion (Fig. 2B) and TCR, HLA-A, HLA-B, and CIITA knockout (Fig. 2C-F), were efficient in the LNP_ortho_ group and comparable to other approaches. TCR knockout was found to be ∼97 to 98% in the LNP_ortho_ group (Fig. 2C). Additionally, to further mitigate GvHD risk, TCR depletion can be applied on engineered cells to achieve >99% TCR removal (Sup. Fig. 2C). Lastly, amplicon-sequencing confirmed that the protein knockouts in the LNP_ortho_ condition were a result of C>T conversions leading to introduction of stop-codons or disrupting splice-sites (Sup. Fig. 2D).

The potential induction of chromosomal translocations across the intended target loci was assessed by ddPCR assays (Fig. 2G). The highest translocation rates (ranging from 0.04% to 0.58%) were observed in the EP_sim_ edited cells. Sequential editing in the EP_seq_ and LNP_seq_ groups led to a reduction in translocations for the temporally distanced edits (i.e. HLA-A/CIITA), but cumulative translocation frequency was still significantly above background levels (Fig. 2G, Sup Fig. 2E). While these results cannot be directly compared to the LNP_ortho_ edited cells (due to differences in primer/probe pairs necessitated by the different sgRNA target sites of Nme2 vs SpyCas9 sgRNAs), we observed only background level of translocation frequencies in the LNP_ortho_ group across all targets with the exception of TRAC/HLA-A (0.13%, Fig.2G). As a positive control for these assays, donor-matched T cells were also produced using an Nme2Cas9 cleavase editor, which in contrast to LNP_ortho_ edited cells, demonstrated significant increases in translocation frequencies beyond background (Fig. 2G). Notably, the HLA-A/HLA-B translocation rates which were determined to be at background levels in LNP_ortho_ edited cells were especially elevated in the Nme2Cas9 cleavase group (3-4%, Fig. 2G), likely due to their close proximity on Chromosome 6p. At present, it is not clear what risk such target-to-target translocations pose, as a separate study investigating multiplex *TRAC, TRBC, and PDCD1* CRISPR/Cas9 cleavase engineered TCR-T cells found that translocation frequency decreased over time in patients. ^32^ Nonetheless, strategies to minimize potential abnormalities are likely more critical as the number of edits introduced increases with cell product complexity. In addition, careful selection of sgRNAs for minimal off-target editing, particularly with cleavase approaches, is equally important as those can further increase translocation rates.

Next, we sought to evaluate the potential for large-scale chromosomal structural re- arrangements such as deletions and truncations, as well as potential translocations from target sites to non-targeted chromosomes. For this, metaphase spreads of allogeneic T cells were characterized using dGH-FISH In-Site assay (KromaTiD) with fluorescent probes bracketing CIITA, HLA-A, HLA-B, and TRAC. Image scoring of 200 metaphase spreads found that LNP_ortho_ cells had background levels (0 events or similar to unedited control cells) of structural re- arrangements across CIITA, HLA-B, and HLA-A (Fig. 2H). In contrast, the cells edited by SpyCas9 nuclease (LNP_seq_ or EP_sim_) had increased levels of translocations across these targets. Additionally, we observed a limited amount of truncations and deletions at Chr.6 (HLA-A or HLA-B probe) exclusively in the nuclease edited groups, highlighting the benefit of base-editing when targeting closely-located genes. CAR insertion into the TRAC loci by CRISPR/Cas9 mediated HDR was performed similarly in all groups, and accordingly large deletions at the TRAC edit site were seen across all conditions (mean: 4.5-10.6% vs. 2.5% in unedited controls), which is consistent with recent reports of 5% to 20% chromosome 14 aneuploidy upon TRAC cleavase editing. ^33,34^ This could be minimized by performing CRISPR/Cas9 editing on T cells in a less activated state; ^34^ however, HDR-mediated insertion at the TRAC locus was most efficient in fully activated T cells (data not shown). Thus, a tradeoff may exist, favoring this edit with higher CAR insertion rates since it enables a lower overall T cell dose to be administered and minimizes dosing of non-therapeutically relevant CAR-null cells. Notably, our long-term in vitro cell culture data demonstrated that Allo-edited cells cannot survive without IL-2 support (data not shown), suggesting minimal genotoxicity concerns.

Finally, the allogeneic CD19-CAR-T cells were evaluated for their functional activity. In a serial rechallenge assay with CD19+ Nalm6 tumor cells, we observed robust long-term tumor cell killing by LNP_ortho_ edited cells comparable with EP_sim_ and LNP_seq_ cells. In contrast, EP_seq_ cells which were exposed to two rounds of electroporation (as a means to reduce translocation events) had a significant drop-off in long term Nalm6 cytotoxicity (Fig. 2I), which was correlated with an increased expression of exhaustion markers PD-1, TIM-3, and LAG-3 (Fig 2J.) and decreased proliferative capacity (Sup Fig. 2F).

Cumulatively, we found that LNP delivery of orthogonal CRISPR/Cas9 cleavase and base-editors provides a tractable platform for engineering an allogeneic T cell therapy, providing advantages over EP or LNP delivery of CRISPR/Cas9 nucleases in terms of improved manufacturability, decreased genotoxicity risk, and maintained or improved CAR-T cell function and phenotype.

### Addressing GvHD and HvG challenges related to Allo T cell therapies

After developing a robust T cell engineering platform, we first tested whether the TCR KO in Allo T cells can prevent GvHD in immunodeficient mice. After dosing luciferase-expressing WT (wild type, i.e. unedited) T cells with endogenous TCRs or engineered Allo T cells with knocked out TCRs into NOG-IL15 mice, we monitored T cell activation and proliferation by IVIS imaging (Fig. 3A). Upon being exposed to mouse antigens for ∼ 20 days, WT T cells began to actively proliferate until they reached a plateau on Day 55 post T cell injection. On Day 62, all mice dosed with WT T cells had clinical signs of GvHD and had to be euthanized. In sharp contrast, Allo T cells with minimal endogenous TCRs (>95% KO) showed no proliferation over time, and no signs of GvHD were detected within 90 days in this study (Fig. 3B). This demonstrates that Allo T cells with minimal endogenous TCRs effectively prevent GvHD.

**Fig. 3.**
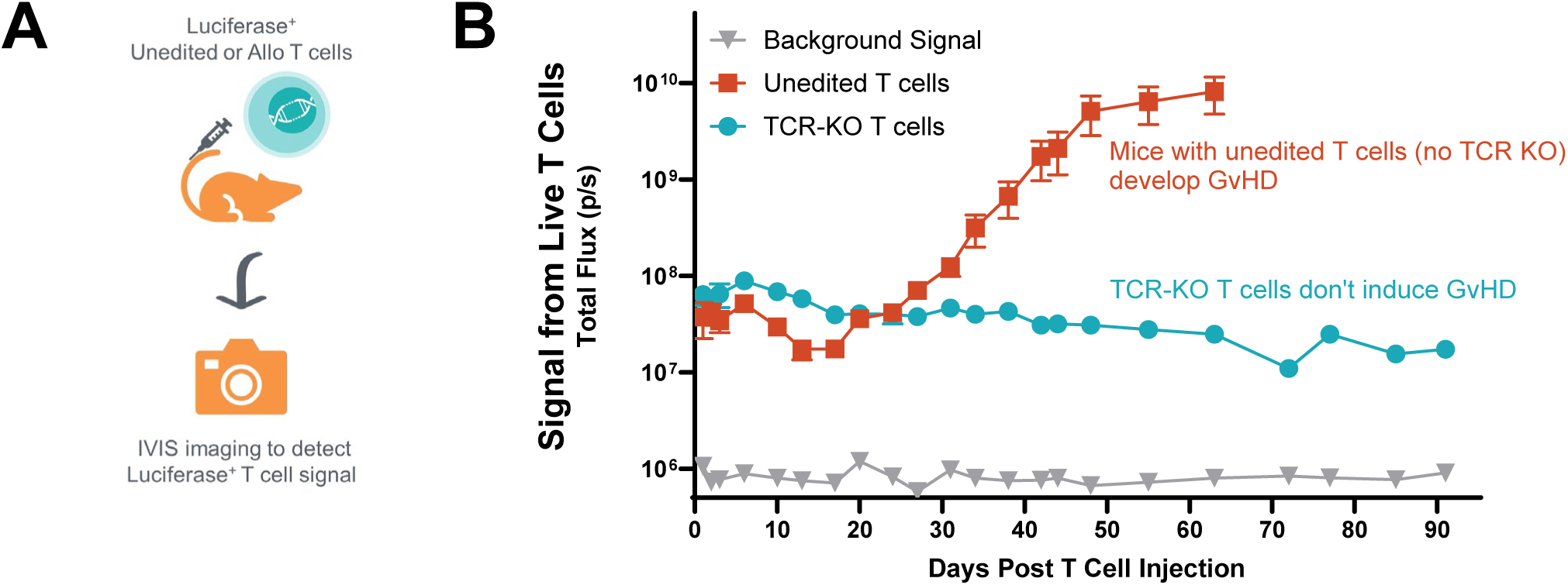
Mice engrafted with TCR-KO T cells did not develop GvHD. A. Schematic of *in vivo* GvHD study workflow. B. Luciferase-expressing unedited T cells or engineered TCR KO T cells were injected into NOG-IL15 immunodeficient mice, and *in vivo* T cell activation and proliferation were monitored by IVIS imaging over 90 days and determined by total flux (photons/s). HBSS buffer was injected in mice as negative control (Background Signal).

Next, we tested the alloreactivity mediated by host T cells. Three groups of donor T cells with different edits (*TRAC* KO alone [TCR KO], *B2M/CIITA/TRAC* KO [B2M KO], and *HLA-A/HLA-B/CIITA/TRAC KO* [Allo T]), were generated. These donor T cells were co-cultured with primed PBMCs from 3 different hosts (autologous host, HLA-A/B/HLA-II mismatched/HLA-C matched, and fully HLA-I/II mis-matched) and evaluated for donor T cell lysis (Fig. 4A).

**Fig. 4.**
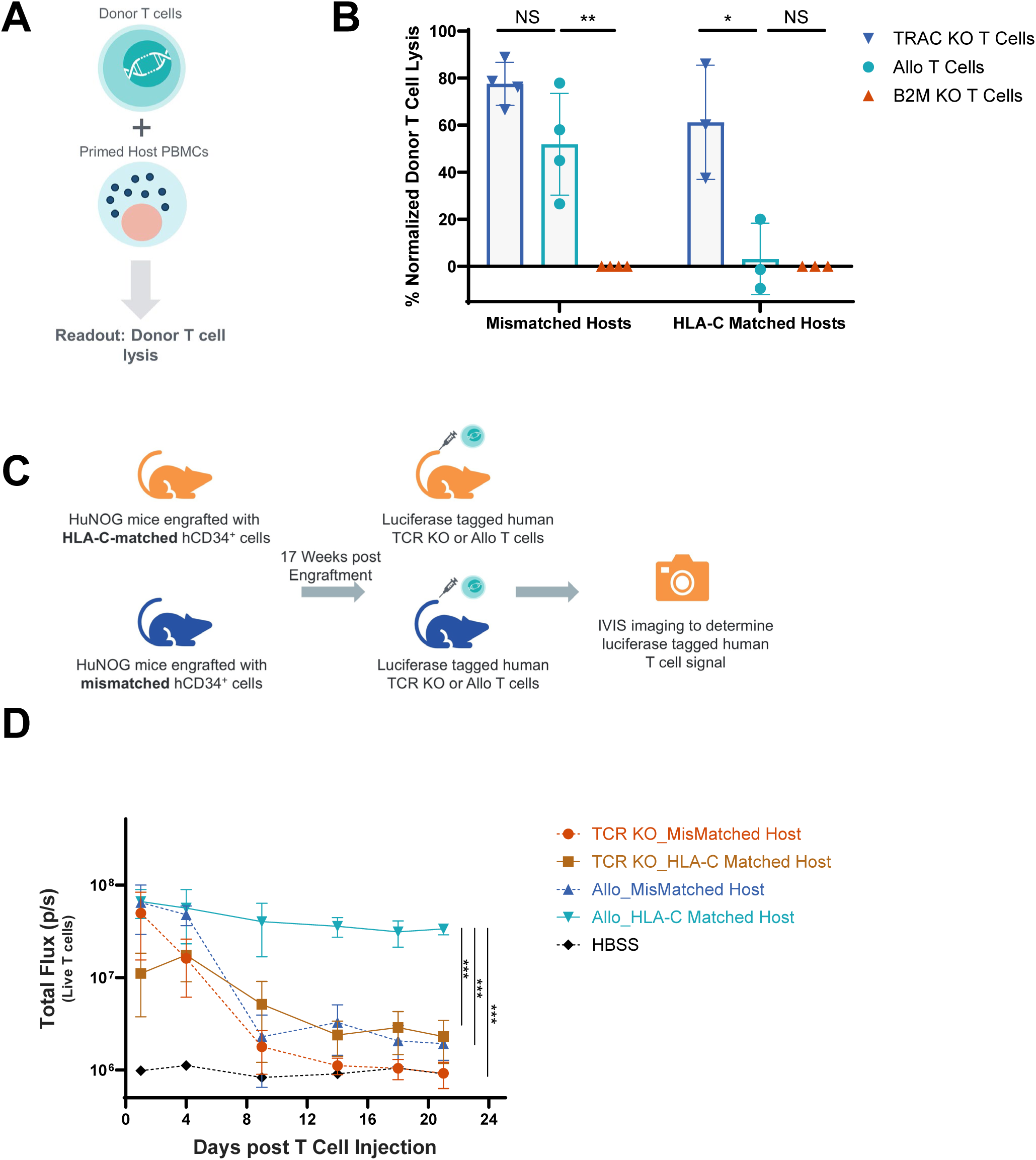
Allo T cells were not rejected by host T cells matched for HLA-C. A. Schematic of *in vitro* host T cell rejection assay. B. TRAC KO T cells, Allo T cells with four Allo edits, and B2M/CIITA/TCR KO (B2M KO) T cells were engineered and co-cultured with primed PBMCs from 3 different hosts, which were autologous host, HLA-C matched host, and mismatched host, at E: T = 3:1. T cell lysis data were normalized based on the data from the autologous host group. Each dot represents a donor/host pair. \**p*<0.05, ** *p*<0.01. NS: not statistically significant. C. Schematic of *in vivo* host T cell rejection model. D. Luciferase-expressing TCR KO T cells (TCR KO) or Allo T cells knocked out for TCR, HLA-A/B and HLA class II (Allo) were injected into humanized NOG mice engrafted with either HLA-C matched CD34+ cells (HLA-C Matched Host) or mismatched CD34+ cells (Mismatched Host). Donor T cell signal was determined by bioluminescent imaging. *** *p*<0.001

Autologous host PBMCs did not kill either TCR KO, B2M KO or Allo T cells, whereas PBMCs from mismatched hosts showed cytotoxicity to both TCR KO T cells and Allo T cells while sparing the B2M KO T cells lacking HLA-I and -II, used as negative controls. Consistent with our hypothesis, HLA-C matched host PBMCs killed TCR knockout T cells, but not Allo T cells (Fig. 4B). To further test this hypothesis, we conducted an *in vivo* study to test the persistence of Allo T cells in the host T cell rejection model. Humanized NOG (huNOG) mice were generated by engrafting human CD34+ cells from a mismatched host or HLA-C matched host into NOG mice. After confirming successful human T cell engraftment in those mice (Sup Fig. 3), we injected luciferized Allo T cells or TCR KO T cells into these huNOG mice and measured T cell signal by IVIS imaging (Fig. 4C). Consistent with *in vitro* data, Allo T cells persisted in mice engrafted with HLA-C matched CD34+ cells (no matching in HLA-A, -B or HLA-II alleles), but not in mice engrafted with fully mismatched CD34 cells (Fig. 4D), indicating that a mismatch in a single HLA allele can be still sufficient to induce rejection, albeit at a slower initial rate than in fully mismatched TCR KO T cells. Control TCR KO T cells were rejected by host T cells regardless of HLA-C matching status, although the single HLA-C match slightly prolonged the persistence of control TCR KO T cells (not statistically significant difference). These findings suggested that Allo T cells with HLA-A/B/class II KO were protected from host T cell attack, and that full protection required HLA-C matching with the host.

As a next step, we interrogated the effect of our Allo approach on NK cell rejection. Similar to the experiments above, we co-cultured Allo T cells and control B2M KO T cells with host NK cells and measured donor T cell lysis in vitro (Fig. 5A). We first evaluated two reported allogeneic approaches, B2M knockout and B2M knockout with HLA-E overexpression. ^26,35^ As shown in Fig. 5B, while wild-type T cells (WT) showed no detectable lysis, even at an effector-to-target (E:T) ratio of 10:1, B2M knockout T cells (B2M KO) exhibited significant lysis in an NK cell dose-dependent manner, attributable to their lack of HLA class I expression. Upon overexpression of HLA-E in B2M knockout T cells (B2M KO+HLA-E), we observed partial protection against NK cell-mediated lysis. However, the majority of these cells were still eliminated by NK cells at higher NK:T cell ratios, suggesting HLA-E overexpression cannot completely suppress NK activity. We compared our Allo approach to the B2M knockout approach with multiple donor/host pairs at an E:T ratio of 10:1. B2M KO T cells were highly susceptible to NK cell mediated killing, ranging from 30-90% depending on the donor/host. In contrast, Allo T cells were significantly more protected from NK cell killing, particularly when T and NK cells were matched for HLA-C (about a 6-fold higher protection (58% vs. 10% killing) vs. only about 3-fold protection with HLA-C mismatched NK cells (60% vs. 22% killing), Fig. 5C-D), indicating that mismatched HLA-C can lead to higher NK cell activation, further favoring HLA-C matching. Next, we tested the persistence of Allo T cells tagged with luciferase in vivo in NOG- IL15 mice engrafted with HLA-C matched NK cells (Fig. 5E). As shown in Fig. 5F, ∼90% of B2M KO T cells were rejected by NK cells engrafted in mice (Sup Fig. 4). Consistent with our *in vitro* data, no significant T cell signal loss was observed in the Allo T cell group (Fig. 5G), indicating that Allo T cells were protected from NK cell-mediated killing *in vivo*.

**Fig. 5.**
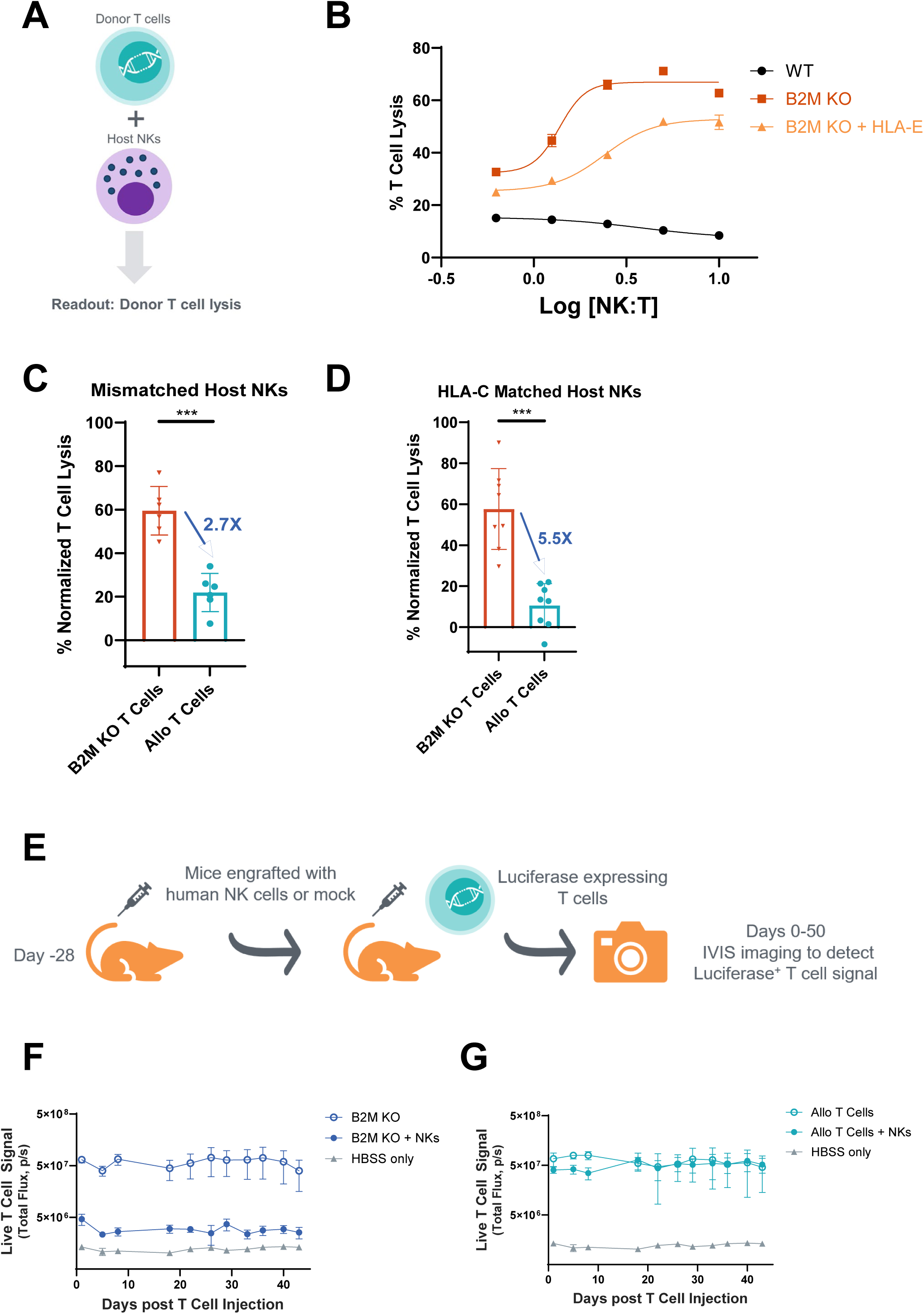
Allo T cells with HLA-C matching were protected from NK killing. A. Schematic of *in vitro* host NK cytotoxicity assay. B. Wild-type T cells (WT), B2M knockout T cells lacking HLA class I (B2M KO) or B2M knockout T cells with HLA-E overexpression (B2M KO + HLA-E) were incubated with different amount of allogeneic host NK cells, and T cell lysis was measured. C & D. TRAC KO T cells, Allo T cells with four Allo edits (Allo T Cells), and B2M/CIITA/TCR KO (B2M KO) T cells were engineered and co-cultured with either HLA-C matched or mismatched NK cells at E:T ratio = 10:1. T cell lysis was normalized based on TRAC KO T cells from the same donor. Each dot represents a donor/host pair. *** *p*<0.001. E. Schematic of *in vivo* host NK cytotoxicity model. F & G. Luciferase-expressing T cells knocked out for B2M/CIITA/TCR (B2M KO) or Allo T cells with four Allo edits (Allo T Cells) were injected into either mock NOG-IL15 mice or NOG-IL15 mice engrafted with HLA-C matched NK cells. T cell signal was determined by bioluminescent imaging.

### Functionality of Allo-TCR-T cells and Allo-CAR-T cells

After we demonstrated Allo T cells had minimal rejection from host T cells and NK cells when matched for HLA-C, we asked whether these Allo T cells were fully functional in the CAR- T or TCR-T cell setting. We first generated control TCR KO cells by CRISPR/Cas9 editing of *TRAC* and inserting either an anti-CD30 CAR or a Wilms’ tumor gene 1 (WT1) specific TCR. For Allo-TCR T cells or Allo-CAR T cells, the three Allo edits (HLA-A, HLA-B, and CIITA) were included during the engineering process. Additionally, for WT1-TCR T cell groups, TRBC knockout was included to reduce TCR mispairing and promote high WT1-TCR expression. ^36^ Allo-T cells were compared with non-Allo edited counterparts for their *in vitro* tumor cell killing capability (Fig. 6A). Both Allo-TCR and Allo-CAR T cells demonstrated potent tumor cell killing *in vitro*, with no significant differences between Allo T cells and their counterparts without the Allo edits (Fig. 6B-C). Moreover, we compared Allo CD30 CAR-T cells and CD30 CAR-T cells with TCR KO by evaluating their *in vivo* tumor clearance capability (Fig. 6D). Again, comparable anti-tumor activity to TCR-KO counterparts was observed *in vivo* (Fig. 6E). The ability of these Allo T cells to effectively kill tumor cells both *in vitro* and *in vivo*, comparable to their TCR-KO counterparts, suggested that engineered Allo T cells with multiple Allo edits maintained their full functionality.

**Fig. 6.**
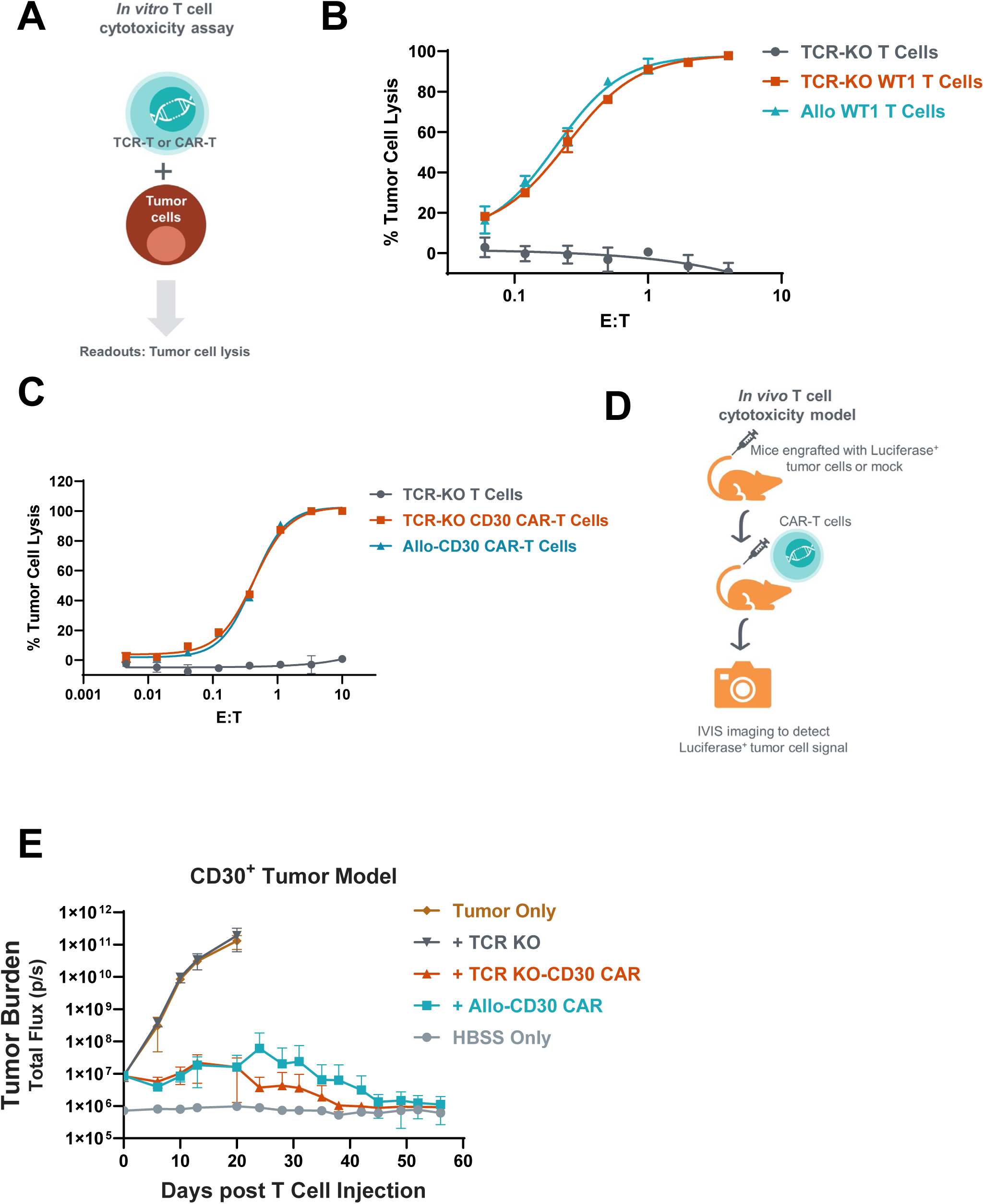
Comparable anti-tumor activity between Allo TCR-T and CAR-T cells and autologous counterparts. A. Schematic of *in vitro* CAR-T or TCR-T cell cytotoxicity assay. B. WT1 TCR-T cells with either TRAC KO only (TCR-KO WT1 T Cells) or Allo edits (Allo WT1 T Cells) were co-cultured with 697 tumor cells at different E:T ratios, and the percentage of tumor cell killing was quantified. TCR KO T cells without WT1-TCR (TCR-KO T Cells) were served as negative control cells. C. CD30 CAR-T cells with either TRAC KO only (TCR-KO CD30 CAR-T Cells) or Allo edits (Allo-CD30 CAR-T Cells) were co-cultured with HH tumor cells at different E:T ratios, and the percentage of tumor cell killing was quantified. TCR KO T cells without expressing CD30 CAR (TCR-KO T Cells) served as negative control cells. D. Schematic of *in vivo* CD30+ tumor suppression model. E. After HH-GFP-Luc2 tumor cells were engrafted into NOG mice, TCR KO T cells without expressing CD30 CAR (TCR KO), CD30 CAR-T cells with TRAC KO only (TCR KO-CD30 CAR) or CD30 CAR-T cells with Allo edits (Allo-CD30 CAR) were injected into tumor-bearing NOG mice. HBSS buffer was injected in mice (HBSS only) as negative control. Tumor burden was monitored twice a week by IVIS imaging.

### Application of the Allo platform in iPSCs

To explore the adaptability of our allogeneic platform beyond T cells, we focused on iPSCs due to their remarkable potential for generating diverse specialized cell types for therapeutic applications. ^37^ We first reprogrammed PBMCs into iPSCs and selected an iPSC clone meeting various quality control criteria, including high post-thaw viability (>90%), normal karyotype, and no mycoplasma contamination (data not shown). Flow data demonstrated that more than 80% of wild type (WT) iPSCs from the selected clone were positive for pluripotency markers SSEA4 and TRA-1-60 (Fig. 7A) and OCT4 and SOX2 (Fig. 7B). Bulk triple KO (TKO) iPSC cells were generated by knocking out *CIITA*, *HLA-A*, and *HLA-B* genes using LNP-delivered CRISPR guides, and a TKO clone was selected with high post-thaw viability (>90%), normal karyotype, and no mycoplasma contamination (data not shown). HLA flow analysis confirmed no HLA-B and HLA-A expression in the selected TKO iPSC clone (Fig. 7C) and ddPCR data substantiated no CIITA gene expression in TKO iPSC cells (Sup Fig. 5A). Tracking of indels by decomposition (TIDE) sequencing analysis also confirmed knock out of *CIITA*, *HLA-A*, and *HLA-B* in the selected TKO iPSC clone (data not shown). Further, TKO iPSC cells expressed similar levels of pluripotency surface markers as WT iPSC cells (Sup Fig. 5B), suggesting TKO iPSC cells maintained their differentiation potential.

**Fig. 7.**
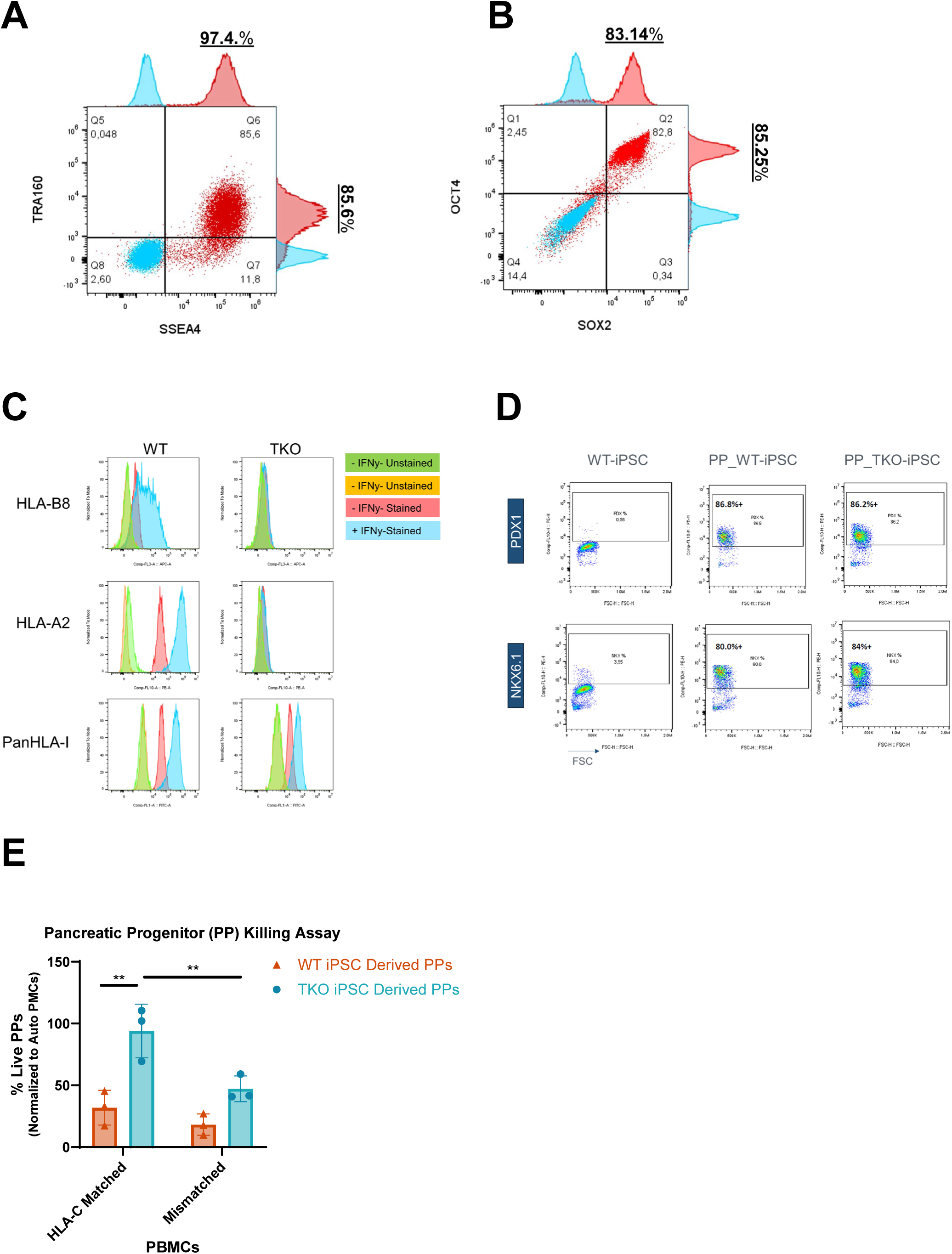
Pancreatic progenitors derived from TKO iPSCs resistant to HLA-C matched PBMC mediated rejection. A-B. After iPSCs were reprogramed from PBMC cells, the expression of pluripotency markers, including SSEA4 and TRA-1-60 (A, red) and OCT4 and SOX2 (B, red), were examined by flow cytometry. Samples stained with isotype control antibodies are shown in blue. C. After WT or TKO iPSC cells were pretreated with or without IFN-γ, surface expression of HLA-B8, HLA-A2, and pan HLA class I were examined by flow cytometry to confirm successful KO of HLA-A and HLA-B. D. After pancreatic progenitor cells were differentiated from WT or TKO iPSCs, those cells were stained with antibodies to detect PDX1 and NKX6.1, two pancreatic β cell markers, to confirm pancreatic progenitor cell differentiation. E. Pancreatic progenitor cells differentiated from WT or TKO iPSCs were co-cultured with primed PBMCs from three different hosts, which were autologous host, HLA-C matched host, and mismatched host, at E:T ratio = 1:1. Pancreatic progenitor cell lysis data were normalized to data from the autologous host group. Each dot represents one PBMC host. ** *p*<0.01.

By leveraging the differentiation potential of iPSCs, we differentiated pancreatic progenitor cells or cardiomyocytes from iPSCs. As shown in Fig. 7D, pancreatic progenitor cells derived from WT or TKO iPSCs showed similar expression levels of pancreatic and duodenal homeobox 1 (PDX1) and NK6 homeobox 1 (NKX6.1), two key transcription factors that are essential for the development and function of pancreatic β cells. Compared to cardiomyocytes derived from WT iPSCs, cardiomyocytes derived from TKO iPSCs had a slightly higher level (91.9% vs 81.7%) of cardiac troponin T (cTNT), a cardiac-specific marker (Sup Fig. 5C). These phenotypic profiling data from pancreatic progenitor cells and cardiomyocytes suggested that triple Allo edits in iPSC cells had minimal impact in iPSC’s differentiation potential. Using these differentiated pancreatic progenitor cells and cardiomyocytes, we performed the mixed lymphocyte reaction (MLR) assay to address the key alloreactivity question. However, very weak alloreactivity was detected with WT iPSC-derived cells, even after pretreatment with IFN-γ to boost HLA expression (data not shown). To solve this assay sensitivity challenge, we first optimized this assay by priming host PBMCs with graft PBMCs, which were used to generate the iPSCs (Sup Fig. 5D). In addition, WT and TKO iPSCs were transduced with lentivirus expressing GFP-luciferase to develop a more sensitive assay tracking survival of luciferized cells. GFP+ iPSC cells were sorted to ensure similar GFP/luciferase expression level (Sup Fig. 5E) and pluripotency markers (Sup Fig. 5F) in WT and TKO iPSCs. In this assay format, we were able to detect significant alloreactivity toward pancreatic progenitor cells derived from WT iPSC cells, while pancreatic progenitor cells differentiated from TKO iPSC were well protected from killing mediated by primed PBMCs from 3 HLA-C matched hosts (Fig. 7E). Consistent with T cell data (Fig. 4B & 4D), greater protection from TKO iPSC differentiated pancreatic progenitor cells were observed when they were co-cultured with primed PBMCs from HLA-C matched vs. mismatched hosts, highlighting the importance of HLA-C matching in suppressing alloreactivity. Similar data were observed in iPSC derived cardiomyocytes (Sup Fig. 5G). These findings suggest that this allogeneic platform can be successfully applied to iPSCs and their differentiated cell types, potentially broadening the scope of allogeneic cell therapies in regenerative medicine.

## Discussion

T cell therapies, including CAR-T, TCR-T, and TIL therapy, have made tremendous progress over the past decade and show potential to revolutionize cancer and autoimmunity treatment. ^1,2,19,24,38,39^ However, lengthy manufacturing times, high costs of goods sold (COGS), and commercial hurdles associated with autologous T cells have significantly impeded their widespread adoption. Aiming to bring the benefits of T cell therapies to eligible patients faster and more cost-effectively, allogeneic T cell therapies have emerged as a potential solution to overcome many of these limitations of autologous T cell therapies. However, persistence of these approaches is still a challenge, affecting their long-term efficacy. ^40^

From experience in hematopoietic stem cell transplantation (HSCT), the primary safety concern with allogeneic T cell therapies is their potential to cause life-threatening GvHD. ^41^ Depletion of TCRαβ+ T cells in allogeneic HSCT is associated with lower incidence and severity of GvHD compared to the incidence in subjects receiving non-depleted transplants. For patients undergoing myeloablative conditioning regimens, HSCT with CD34+ cells with T cell depletion was associated with a low rate of mostly grade 1 or 2 acute and chronic GvHD, containing a maximum dose of αβ T cells of 1 × 10^5^/kg. ^42^ Recent clinical trials of allogeneic CAR T cell therapies also have highlighted the risk of acute graft-versus-host disease (aGvHD) associated with residual TCRs in donor-derived T cells. In a trial using allogeneic CD19 CAR T cells for relapsed/refractory B-cell precursor acute lymphoblastic leukemia, one out of thirteen patients developed aGvHD. ^43^ More strikingly, in a trial of allogeneic GD2 CAR T cells for relapsed or refractory neuroblastoma, four out of five patients experienced aGvHD. ^6^ Both trials utilized TCR-positive T cells from either fully HLA-matched or haploidentical donors. The higher incidence of aGvHD in the GD2 CAR T cell trial particularly underscores the potential risks associated with residual TCRs in more persistent allogeneic T cell products, emphasizing the need for strategies to mitigate this complication in future allogeneic CAR T cell therapies. Two major approaches to address this issue are the use of virus-specific memory T cells or the knockout TCRs in T cells. Tabelecleucel, an allogeneic Epstein-Barr virus (EBV)-specific T-cell immunotherapy, ^24^ and EBV-specific allogeneic CD30 CAR-T therapy ^25^ exemplify the first approach. The caveat is the generation of EBV-reactive T cells requires multiple rounds of stimulation to enrich those T cells, potentially leading to T cell exhaustion and an intrinsically short-lived phenotype ^44^. Given that the root cause of GvHD is the presence of alloreactive TCRs in the allogeneic T cells, another popular method is to engineer T cells with minimal endogenous TCR expression, which has been applied in many T cell products via various gene editing modalities, including transcription activator-like effector nuclease (TALEN), CRISPR Cas9, and base editor. ^26,45,46^ More recent data from multiple clinical trials demonstrate no detected GvHD in TCR-KO allogeneic T cell therapies, ^26,47–49^ suggesting this issue has been well addressed with either highly efficient KO and/or add-on TCR-positive T cell depletion at the end of the manufacturing process.

The next challenge is mitigating host immune system mediated rejection of allogeneic products. Several strategies have been explored, including intense lymphodepletion (LD) regimen, ^50,51^ immune evasion modifications to render allogeneic T cells invisible to the host immune system, ^26,35^ and armored T cell approaches in which allogeneic T cells are engineered to recognize and eliminate alloreactive immune cells. ^52^

At least two intense lymphodepletion methods have been investigated to improve allogeneic T cell expansion and persistence. The first is an enhanced lymphodepletion chemotherapy regimen (eLD) evaluated in PBCAR0191 T cells. After the enhanced LD treatment, PBCAR0191 CAR-T kinetics were profoundly improved compared to standard LD, with both peak cell expansion and area under the curve increasing 80-fold and increased T cell persistence from 28 days to 90 days. ^50^ Another approach is to leverage the anti-CD52 monoclonal antibody to deplete CD52-expressing immune cells and achieve the prolonged lymphodepletion effect without high chemotherapy doses, while sparing therapeutic T cells by disrupting CD52 expression. For example, ALLO-715 is a genetically modified CAR-T cell therapy with *CD52* gene KO, permitting the use of anti-CD52 monoclonal antibody for selective and transitory host lymphodepletion. In the presence of anti-CD52 monoclonal antibody, ALLO- 715 T cell’s persistence was observed up to Month 4 in Dose Level 3. ^51^ Unfortunately, these intense lymphodepletion approaches have raised some safety concerns pertaining to infections, even leading to some reported cases of non-disease progression deaths due to infections (fungal pneumonia and adenovirus hepatitis) in the PBCAR0191 CAR-T trial ^50^ or the ALLO-715 trial. ^53^ These infectious death cases reported in both trials were most likely associated to intense lymphodepletion.

To address the severe infection risk associated with intense lymphodepletion, immune evasion modifications in T cells as an alternative strategy are being explored. One approach eliminates HLA class I surface expression by knocking out *B2M*. CTX130, an allogeneic CD70- targeting CAR T-cell product, disrupts both *TRAC* and *B2M*. Pharmacokinetic (PK) analysis from the recent Phase 1 trial revealed that, at dose level of 9 × 10^8^ T cells, CTX130 cells were no longer detected in patients’ blood by Day 28 post T cell injection (N=4). ^26^ The poor persistence of *B2M* KO T cells may be attributed to their rapid elimination by host cytotoxic NK cells through the well-characterized “missing-self” mechanism. ^27^ Similarly, the persistence of P-BCMA-ALLO1, an allogeneic CAR-T product candidate with TCR knockout and partial B2M knockout, was up to 40 days. This short persistence was observed even with intense lymphodepletion as pre-conditioning, which included high dose of cyclophosphamide (750 mg/m²/day) and fludarabine (30 mg/m²/day). ^54^

To protect allogeneic T cells from both host CD8 and NK cell attack, another strategy involves expressing HLA-E, an NK inhibitor, in T cells deficient for TCR and HLA Class I. ^55^

While promising, this approach has caveats due to HLA-E’s ability to bind both inhibitory receptor NKG2A and activating receptor NKG2C on NK cells, leading to varying protection based on the relative expression of NKG2A vs NKG2C on host NK cells. ^56^ This was also confirmed in our studies, showing high variability and incomplete protection using this approach in cell-based assays. Clinical data testing this approach are still pending.

Another approach to inhibit host CD8 and NK cells is to express CD47 in T cells. A recent study reported that CD47 overexpression can protect HLA class I and II-disrupted CAR T cells against host NK cells. ^57^ However, clinical data with engineered T cells are pending as well. UP421, an allogeneic primary islet cell therapy engineered with the same technology, was reported with early but promising results. Specifically, four weeks after UP421 islet cell transplantation in the first treated Type 1 Diabetes patient without immunosuppression, there was evidence of pancreatic beta cell survival and function, as demonstrated by the presence of circulating C-peptide.

Besides downregulation of MHC class I, tumor cells or virus-infected cells can evade immune eradication and become “invisible” by losing adhesion molecule CD54 and CD58, which contribute to synapse formation between immune cells and target cells. With this observation, a recent study demonstrated that simultaneous deletion of CD54 and CD58 on *B2M* KO T cells could limit NK cell activation universally and partially protect *B2M* KO target cells from host immune cell rejection. ^58^ It will be interesting to see how this preclinical study translates in a clinical setting.

Recently, using an *in vivo* genome-wide CRISPR KO screen, one group discovered that deletion of either *Fas* or *B2M* in Allo T cells enhanced their survival in immunocompetent mice. ^59^ Unlike *B2M* KO T cells, *Fas* KO T cells maintain normal expression levels of HLA Class I and demonstrated resistance to NK cells in both *in vitro* and *in vivo* studies. Based on these findings, the authors concluded that *Fas* ablation represents a more robust and effective target than *B2M* for developing off-the-shelf allogeneic T cell therapies. However, the authors also pointed out that Fas plays a critical role in maintaining immune homeostasis and preventing autoimmunity. Consequently, its ablation in CAR T cells could raise safety and regulatory concerns that need to be carefully addressed.

Yet another approach enables allogeneic T cells to attack and deplete alloreactive host cells. For example, alloimmune defense receptor (ADR) T cells express a CAR that selectively recognizes 4-1BB, a cell surface receptor temporarily upregulated by activated (alloreactive) T cells and NK cells ^60^. However, this approach may have several limitations. First, other immune cells, including activated B cells, follicular dendritic cells, follicular helper T cells monocytes, eosinophils and mast cells, can also express 4-1BB transiently. Future studies should evaluate whether ADR T cells broadly deplete immune cells and suppress immune response induction when needed. Secondly, it is also possible that ADR activity may result in a temporary suppression of productive systemic T-cell responses in the tumor or pathogen specific settings, leading to similar side effects as seen with enhanced lymphodepleting chemotherapy or anti- CD52 antibody. Lastly, as observed by the authors, the expression of 4-1BB on activated ADR T cells could cause unfavorable fratricide issue. ^60^ Another armored Allo approach leverages a CD70 CAR insertion to deplete activated, alloreactive lymphocytes that upregulate CD70. ^61^ Similar to 4-1BB, CD70 is expressed on regulatory T cells (Tregs) and various activated immune cells, including T cells, B cells, dendritic cells and NK cells. ^62,63^ Due to CD70’s broader expression profile, CD70 CAR-based allogeneic approach may inadvertently suppress normal defensive immune responses.

Different from the approaches alluded to above, our allogeneic strategy is to facilitate full matching of the product to the host, knocking out *HLA-A*, *HLA-B*, and *CIITA* genes and matching one residual HLA-C allele. To enable this multiplexed editing strategy, we developed a differentiated and robust T cell engineering process. First, with our deep expertise in LNP biology, ^64,65^ we developed a T cell specific LNP delivery process, which leads to high editing efficiency, higher T cell viability, more robust cell expansion, and much lower genomic alterations compared to electroporation. LNPs also enable multiplex sequential editing i.e. edits on different days, which offers more editing flexibility and minimizes competition of edits and potential genotoxicity concerns. Secondly, the development of a high-fidelity Nme2Cas9 base editor enables multiplex knockout, which leads to minimized indels, DNA double strand breaks, and translocations. Thirdly, with CRISPR/Cas9 cleavase, site-specific insertion of an AAV-based delivery of an HDR template encoding a CAR or therapeutic TCR is achieved to minimize the risk of insertional mutagenesis and cell transformation. These risks have been observed in some cases related to random integration of retroviral or lentiviral vectors that may disrupt tumor suppressor genes or activate oncogenes. ^66–68^

Our *in vitro* and *in vivo* data demonstrated that highly efficient TCR knockout (>95%) eliminated GvHD concerns, and our allogeneic T cells were protected from both host T cell and NK cell mediated killing. Because of the low immunogenicity of these engineered T cells, this approach is expected to be compatible with standard lymphodepletion, rather than relying on enhanced lymphodepletion. Importantly, Allo TCR-T and CAR-T cells showed comparable anti-tumor activity to autologous counterparts in both *in vitro* and *in vivo* studies, suggesting this allogeneic platform is readily deployable for TCR-T and CAR-T cell therapies.

Our preclinical persistence data is further supported by subsequent clinical data from multiple studies. The first study is the CB-010 ANTLER Phase 1 trial presented at 2024 ASCO. ^47^ CB-010 is an allogeneic anti-CD19 CAR-T cell therapy derived from healthy donor T cells with 3 edits, including knockout of *TRAC* and *PD-1* and insertion of a CD19-specific CAR into the *TRAC* locus. A retrospective analysis which stratified patients on the basis of HLA matches to donor T cells found a median progression free survival (PFS) of 14.4 months in patients treated with CB-010 with ≥4 HLA matches (N=13), compared to just 2.8 months for patients treated with CB-010 with ≤3 HLA matches (N=33), suggesting that partial HLA matching with ≥4 matching HLA alleles may improve durability of response to CB-010 Therapy. This observation is aligned with PK data, which showed that higher numbers of matched HLA alleles between the CB-010 donor and recipient patient correlated with increased CAR-T cell expansion and persistence compared to lower numbers of matched HLA alleles. ^47^ Because they did not knock out any HLA alleles, they predicted at least 13 manufacturing batches will be needed to cover 90% US patients with ≥4 matched alleles. ^47^ The previous data from the hematopoietic stem cell transplantation field suggest that 4-allele matching is not ideal to address the host immune cell mediated rejection. The data presented here clearly show that mismatches even in a single allele can lead to significant rejection by T and NK cells. In addition, the ambiguity of more than 4-allele matching may result in potential variability of CB-010’s clinical performance across patients based on total levels of HLA-matching. Furthermore, we determined that only 7 batches will be able to cover about 90% of US patients with full match. In another study, the authors tested TyU19 T cells in three patients with refractory autoimmune diseases. TyU19 is a healthy donor-derived allogeneic CD19-targeting CAR-T product with 5 knockouts, including *TRAC*, *HLA-A*, *HLA-B*, *CIITA* and *PD-1*. ^49^ Despite the small sample size (N=3) and short follow-up time (up to 6 months), this group reported encouraging efficacy and over-3-month allogeneic T cell persistence data, providing clinical proof of concept for our allogeneic platform. However, different from our Cas9 base editor / cleavase and LNP delivery, they performed 5 knockouts by Cas9 cleavase and electroporation delivery, leading to approximately 1%–4% translocation events and raising an increased safety-related genotoxicity concern.

Human pluripotent stem cells (hPSCs) derived cell therapies, particularly those using iPSCs, hold immense promise for regenerative medicine, offering a path to create customized cell therapies for various diseases, including degenerative ocular diseases, Type 1 diabetes, Parkinson’s disease, and cardiovascular disorders. ^69–72^ As of December 2024, a total of 116 clinical trials have been conducted to test 83 hPSC-derived products for 34 different indications. ^73^ Several early but encouraging clinical trial results have been reported. For example, VX-880, a stem cell-derived islet product, is currently in a clinical trial for Type 1 diabetes. Recent data showed that all six patients who received these cells demonstrated insulin production and improved glycemic control, including reduced levels of HbA1c and reducing or eliminating the need for exogenous insulin. ^70^ This Phase 1/2 trial for VX-880 has converted to a Phase 1/2/3 pivotal trial in November 2024, enrolling a total of 50 patients.

Because stem cells are renewable cell sources and they can differentiate into various specialized cell types, iPSC-based off-the-shelf approaches may represent the next generation of allogenic cell therapies. Similar to bone marrow transplantation and allogeneic T cell therapies, the holy grail of allogenic iPSC approaches is to achieve graft transplantation without immunosuppression. While the VX-880 clinical trial shows promising results, it still requires systemic immunosuppression and encapsulation of allogeneic cells in a device to protect them from short and long-term immune rejection, which is not ideal due to side effects related to systemic immunosuppression or viability concerns due to encapsulation. Using our approach, we differentiated iPSC cells into pancreatic progenitor cells or cardiomyocytes and demonstrated that both cell types derived from HLA-A/B/CIITA TKO iPSCs were protected from host T cell mediated rejection in an HLA-C matching dependent manner.

In conclusion, we have developed a differentiated and durable allogeneic T cell strategy by editing *TRAC*, *HLA-A*, *HLA-B*, and *CIITA* genes, and established a robust T cell engineering platform by leveraging orthogonal Cas9 enzymes and an AAV/LNP delivery system. Our preclinical studies have generated promising *in vitro* and *in vivo* data to support first-in-human clinical study. Moreover, our data demonstrate the potential of this platform in advancing iPSC-derived regenerative therapies.

## Materials & methods

### T cell engineering

Allogeneic T cells were engineered by either (1) Ribonucleoprotein (RNP) electroporation (EP) of SpyCas9-cleavase RNPs or (2) Lipid nanoparticle (LNP) delivery of SpyCas9-cleavase mRNA and sgRNA or (3) a combination of LNP delivered SpyCas9-cleavase mRNA/sgRNA and Nme2Cas9 base editor mRNA/sgRNA. Engineering of these cells began with enrichment of CD8+ (Miltenyi Biotec, Cat. 130-045-201) and CD4+ (Miltenyi Biotec, Cat. 130-045-101) T cells from three healthy PBMC donors and cryopreservation in CryoStor10 (Stemcell Technologies, Cat. 07930). CD4+ and CD8+ T cells were thawed on Day 0 and combined at a 1:1 ratio and rested overnight in T cell activation media (TCAM: CTS OpTmizer T cell Expansion SFM (Gibco, Cat. A3705001) supplemented with 2.5% (v/v) of GemCell Plus Human AB Serum (Gemini, H36X00K), 1x Glutamax (Gibco, Cat. 35050-061), 10 mM HEPES buffer, 1% of Penicillin-Streptomycin, and 20 ng/mL of recombinant human interleukin-2 (Peprotech, Cat. 200-02), 5 ng/mL of recombinant human interleukin-7 (Peprotech, Cat. 200-07), and 5 ng/mL of recombinant human interleukin-15 (Peprotech, Cat. 200-15). The following day (Day 1), cells were activated with TransACT^TM^ (1:100 dilution, Miltenyi Biotec, Cat. 130-111-160) and the gene edits for CIITA and HLA-B were performed. Cas9 RNPs were complexed and nucleofected (Lonza) in P3 buffer using the EH115 program as previously described ^36^. For LNP delivery, nanoparticles encapsulating the appropriate mRNA and sgRNA were added to the respective cultures supplemented with (10 µg/mL) rApoE3 (Peprotech, Cat. 350-02). On Day 3 the edit for HLA-A was performed along with targeted insertion of the CAR transgenes into the *TRAC* locus using an AAV homology directed repair template as previously described. ^74,75^ To improve HDR-mediated insertion, a small molecule DNA-PK inhibitor was added to the culture to improve the insertion rates ^75^. The cells were incubated for 24 h and then transferred to 6-well GREX plates (Wilson Wolf, Cat. 80240M) for expansion in T cell expansion media (TCEM: CTS OpTmizer media supplemented with 5% v/v CTS Immune Cell Serum Replacement (Gibco, Cat.

A2596101), 1x GlutaMAX, 10 mM HEPES, 100 U/mL PenStrep, 20 ng/mL IL-2, 5 ng/mL IL-7, and 5 ng/mL IL-15). Cells were harvested on Day 11 or Day 15 and cryopreserved for future use. Where specified residual TCR+ cells were depleted using TCRα/β microbeads (Miltenyi Biotec, Cat#200-070-407) following the manufacturers’ recommendations.

### Engineered T cell characterization

The engineered cells were characterized for gene editing and insertion rates by flow cytometry (Cytoflex LX, Beckman Coulter) using antibodies specific to CD3 (BioLegend, Cat. 317336), CD4 (BioLegend, Cat. 317434), CD8 (BioLegend, Cat. 301046), HLA-A (BioLegend, Cat. 343320 and eBiosciences Cat. 17-5754-42), HLA-B (Miltenyi Biotec, Cat. 130-118-332 and 130-118-366), CIITA (BioLegend, Cat. 361712). Memory phenotype was characterized using antibodies targeting CD62L (BioLegend, Cat. 304810) and CD45RA (BioLegend, Cat. 304126). Editing frequency and type was further confirmed by targeted amplicon-sequencing using primers flanking the corresponding editing site and sequenced using the Illumina NextSeq2000. TCR depletion was also confirmed by flow cytometry (Cytoflex LX, Beckman Coulter) using antibodies specific to CD3 (BioLegend, Cat. 300430), TCRα/β (Miltenyi Biotec, Cat. 130-113-530), TCR γ/δ (BD Pharmingen, Cat. 555718), and viability dye ViaKrome (Immunotech, Cat.C36628).

### Prediction of HLA-C type coverage in US population

The coverage of HLA-C types in the US population was evaluated using data from 2.1 million donors in the BeTheMatch database. The analysis, performed using R, utilized packages such as dplyr, tidyr, reshape2, and ggplot2. The cumulative percentage for a specific HLA-C type was determined by summing its percentage and those of more frequent HLA-C types. This methodology was also applied to analyze HLA-C coverage across different racial subgroups, allowing for a more granular analysis of HLA-C coverage across different US demographic groups.

### *In vitro* cytotoxicity rechallenge assay

The functionality of allogeneic CAR-T cells engineered by different methods was evaluated in a serial tumor rechallenge format. For this, CD19 CAR T cells were co-cultured with GFP-positive NALM6 cells at an E:T ratio of 1:1, and the GFP loss, i.e., tumor control, was tracked by IncuCyte® live cell imaging. Fresh tumor cells were added to the co-culture every 48 h, and the T cells were subjected to repetitive rechallenge cycles until they lost the ability to kill the tumor cells.

### Genotoxicity studies

The occurrence of chromosomal translocations across target loci was quantified using ddPCR (Bio-Rad). For each potential translocation event, custom primer/probe assays were designed and reaction conditions optimized using synthetic gBlocks. Genomic DNA was extracted from engineered cells using DNeasy blood & tissue kit (Qiagen, Cat. 69506). For ddPCR reactions, droplets were generated in the Bio-Rad Automated Droplet Generator and read on the QX200^TM^ Droplet Reader. To further characterize potential large scale genomic aberrations such as deletions, truncations, and translocations, a directional genomic hybridization (dGH) in-Site^TM^ assay was performed on engineered cells (KromaTiD) using custom fluorescent probe brackets for each edit’s sites.

### *In vivo* GvHD model

Female hIL15-NOG mice were sourced from Taconic Biosciences (Albany, NY), at least 7-8 weeks old, and used in the studies. All mice were maintained under pathogen-free conditions under protocols approved by the institutional guidelines and approval by local authorities. To study GvHD, mice were injected with unedited or TCR knockout T cells. To examine background signal from IVIS machine, HBSS buffer was injected in mice as negative control. Overall health was monitored by clinical observations and weight measurements. IVIS imaging was performed to identify luciferase-positive T cells in mice by IVIS spectrum. Mice were prepared for imaging with an injection of Xenolight D-luciferin (Revvity, Cat. 122799) at 10 µL/g body weight per the manufacturer’s recommendation. Region of Interest (ROI) was specified, and total flux values were calculated using Living Image Software. All the data was plotted and analyzed using GraphPad Prism software.

### *In vitro* T cell rejection assay

Allo edited donor T cells and control T cells were tested with either complete HLA-I mismatched PBMCs or HLA-C matched PBMCs and compared to autologous cells as a negative control.

PBMCs and unedited T cells from the selected donor were thawed in T cell growth media (TCGM) composed of CTS™ OpTmizer™ SFM (Gibco, Cat. A1048501), Human Serum AB (GeminiBio, Cat. 100-512), HEPES (Gibco, Cat. 15630-080), GlutaMAX Supplement (Gibco, Cat. 35050-061), and Penicillin-Streptomycin (Gibco, Cat. 15070-063) and further supplemented with 10 ng/mL of recombinant human IL-2 (Peprotech, Cat. 200-02), 5 ng/mL IL-7 (Peprotech, Cat. 200-07), 5 ng/mL IL-15 (Peprotech, Cat. 200-15). Cells were rested in a 37°C incubator overnight. The following day, unedited donor T cells were irradiated at 5000 rad and co-cultured with CD56 depleted Host PBMCs using CD56 Microbeads (Miltenyi Biotec, Cat. 130-097-042) at an E: T ratio of 1:1 in a 6 well G-Rex in TCGM media containing cytokines. Priming was allowed to take place for 7 days. After priming for 7 days, PBMCs were washed and co-cultured with GFP-Luciferase expressing Allo edited T cells or GFP-Luciferase expressing control T cells at an E: T ratio of 3: 1 for 24 h. Bright Glo Luciferase was added to the plate to determine killing of luciferase expressing donor T cells by primed PBMCs. Percent killing was normalized with B2M KO T cells.

### *In vitro* NK rejection assay

Cryopreserved host HLA-I mismatched or HLA-C matched NKs along with engineered Allo edited donor T cells were thawed and rested overnight in T cell growth media (TCGM) as described above for T cells and 20 ng/mL of recombinant human IL-2 (Peprotech, Cat. 200-02) & 5 ng/mL IL-15 (Peprotech, Cat. 200-15) for NK cells. The following day engineered donor T cells were co-cultured with Cell Trace Violet (CTV) (Invitrogen, Cat. C34557) labelled Host NK cells at an E: T ratio of 10: 1 in a 96-well plate and incubated for 18-20 h at 37°C. The next day, NK cell killing activity was evaluated by performing flow cytometry by staining with the live/dead cell dye DRAQ7 (BioLegend, Cat. 424001) and reading the plate on Cytoflex LX or MACS Quant flow cytometry machines. Live T cells were gated on CTV & DRAQ7 double negative cells. B2M KO and Allo groups were normalized to TRAC KO groups. Average for each duplicate was calculated and reported for mismatched NKs or HLA-C matched NKs.

### *In vitro* CAR-T-mediated killing assay

T cells expressing an anti-CD30 CAR with or without Allo edits were tested for their cytotoxicity against luciferase expressing CD30+ HH tumor cell lines. ^76^ HH tumor cells were thawed and maintained in culture for at least 7 days before setting up the killing assay. Engineered T cells were thawed and rested overnight in pre-warmed T-cell growth media (TCGM) as described above. Killing assay was setup the following day by co-culturing HH tumor cells with T-cells at titrated E:T ratios starting from 10:1 and serially diluting 3-fold over 8 points. Tumor cell killing was allowed to take place for 18-20 h at 37°C. Tumor cell killing was detected by using Bright-Glo™ luciferase assay system (Promega, Cat. E2620). The plate was read for luminescence with a Promega Promax plate reader. The percent killing was calculated from the luminescence by normalizing with the tumor cell only group as 0% killing.

### *In vitro* WT1 TCR-T killing assay

Engineered T cells were thawed and rested overnight in T-cell growth media (TCGM) as described above. On the following day, co-cultures with tumor cell line 697 expressing luciferase and GFP (ACC 42/DMSZ) were established at titrated E:T ratios. To probe real-time kinetics of cytotoxicity, the Incucyte® S3 Live-Cell Analysis Systems was used where loss of fluorescence from GFP-labelled lines quantifies target cell lysis. GFP measurements were normalized to the tumor cell only control (100%) for each timepoint.

### *In vivo* T cell rejection model

Female HuNOG (Model #HSCCB-NOG) mice with the HLA phenotypes suitable for our ‘HLA-C match’ and ‘mismatch’ parameters were sourced from Taconic Biosciences (Albany, NY). These humanized immunodeficient NOG mice were engrafted with human CD34+ cord blood cells and are replete of human B cells and CD3+ T cells and thereby provide stable engraftment of primary human immune cells enabling long term studies. A QC was performed at 12 weeks post cord blood engraftment at Taconic to determine hCD45 cell engraftment levels. Animals with appropriate levels of hCD45 cell engraftment were randomized and assigned to study groups.

Engineered T cells expressing GFP-Luc were thawed and washed with HBSS (Gibco, Cat. 14025126) and prepared at a concentration of 5 x 10e^6^ live GFP-Luc+ cells per mouse and administered intravenously for T cell dosing. Overall animal health was monitored by weight measurements and clinical observations. Bioluminescence based IVIS imaging was performed to track luciferase positive cells in vivo post T cell dosing as described above.

### *In vivo* NK rejection model

Female hIL15-NOG mice sourced from Taconic Biosciences (Albany, NY), at least 7-8 weeks old, were used in these studies. All mice were maintained under pathogen-free conditions under protocols approved by the institutional guidelines and approval by local authorities. Primary human NK cells were isolated from leukopaks (StemCell Technologies) and 1.5 × 10^6^ NK cells, suspended in HBSS (Corning, Cat. 21-022-CV), were injected intravenously in hIL15-NOG mice. Four weeks later, mice were injected with T cells in NK engrafted mice, and non-NK engrafted mice were used as controls. To examine background signal from IVIS machine, HBSS buffer was injected in mice as negative control. Overall health was monitored by clinical observations and weight measurements. IVIS imaging was performed to identify luciferase-positive T cells in mice by IVIS spectrum as described above.

### *In vivo* CD30+ tumor model

Female NOG mice sourced from Taconic Biosciences (Albany, NY), that were at least 7-8 weeks old were used in these studies. Mice were engrafted intravenously with 0.3 x 10^6^ HH-GFP-Luc2 tumor cells. At day 4 post tumor engraftment, mice were randomized based on tumor burden.10 × 10^6^ CAR+ T cells, suspended in HBSS (Corning, Cat. 21-022-CV), were intravenously injected. Overall health was monitored by clinical observations and weight measurements. Tumor burden was monitored twice a week using bioluminescence measurements by IVIS Spectrum as described above. Region of Interest (ROI) was specified over each mouse, and total flux values were calculated using Living Image software.

### iPSC reprogramming

The i001 iPSC line was generated at CCRM/OmniaBio (Toronto, ON, Canada) from unrelated human peripheral blood leukopak donors, obtained from Stem Cell Technologies. PBMC vials were thawed, resuspended in 2% FBS (Life Technologies, Cat. 16000-044) / PBS (Life Technologies, Cat. 12491-015) solution and the CD34+ fraction was isolated using the Human Whole Blood / Buffy Coat CD34+ Selection kit (Stem Cell Technologies, Cat. 18086). The CD34+ fraction was washed in 2% FBS+1mM EDTA solution, then resuspended in StemSpan SFEM II medium (Stem Cell Technologies, Cat. 09655) containing CD34+ Expansion Supplement (ExS, Stem Cell Technologies, Cat. 02691) + 1% pen-strep (Life Technologies, Cat. 15140-122) and cultured for 8 days at 37°C in a normoxic incubator to expand, feeding every other day. On day 8, CD34+ cells were counted and 0.5 × 10⁵ cells were resuspended in fresh SFEM II + ExS medium for reprogramming to iPSCs via Sendai virus. Cells were inoculated with virus from the Cytotune –iPS-2.0 Sendai Reprogramming kit (Life Technologies, Cat. A16517) at an MOI of 5:5:3 (hKOS:hc-Myc:hKlf4) for 24 h at 37°C. On day 2 post-transduction, CD34+ cells were plated at low densities on 6-well plates coated with Biolaminin-521 (Stem Cell Technologies, Cat. 200-0117) in SFEM II + ExS medium, then fed on day 3 with TeSR-E7 medium (Stemcell Technologies, Cat. 5914) + 1% Pen-strep (Life Technologies, Cat. 15140-122) until approximately days 16- 21 post-transduction, when iPSC-like dense cell colonies appear. Multiple colonies were manually picked into individual 6-well plates coated with Matrigel (Corning, Cat. 354277) using 23Gx1 SafetyGlide Needles (BD Biosciences, Cat. 305902) and clones were expanded in mTesR medium (Stem Cell Technologies, Cat. 85851) + 1% Pen-strep. Established colonies were further expanded and matured by clump passaging using Gentle Cell Dissociation Media (GCDR, Stem Cell Technologies, Cat. 100-0485) to P5, then adapted to single cell passaging by incubating with 500uL GCDR for 10 mins at 37C, and reseeding at 1-2 × 10⁵ cells/cm^2^ in mTesr + 10uM ROCK inhibitor (Rocki, Y-27632) (Biotechne- Tocris, Cat. 01214) for at 3 subsequent passages. Finally, single cell-adapted iPSC clones were heat shocked at 39°C for 48 h to remove residual sendai virus, before returning to 37°C for maintenance. At this stage, at least 2 established clones per line were frozen as working banks in mTesR (Stemcell Technologies, Cat. 05855) prior to characterization.

### iPSC characterization

Post-thaw viability of cryopreserved iPSC clones was confirmed at >90% in all lines, and lines were confirmed as mycoplasma-negative using the Lonza MyoAlert Plus Kit (Lonza, Cat. LT07- 710). The genetic identity of iPSC lines was performed by polymorphic short tandem repeat analysis (STR). A 6-well plate well of iPSCs was scraped into a microcentrifuge tube DNA was extracted using QIAMP DNA extraction kit (Qiagen, Cat. 51304), as per manufacturer’s instructions. DNA concentration and quality was assessed using a Nanodrop 2000, and 400 ng of iPSC and parental PBMC gDNA was submitted for independent analysis by The Centre for Applied Genomics (TCAG). PCR profiling of 9 STR regions plus Amelogenin for gender determination confirmed matching of all iPSC lines with parental PBMCs. Structural chromosomal assessment of all iPSC lines was performed via karyotype testing by WiCell, with G-banding analysis detecting structural abnormalities of size >3-10Mb.

### Generation of TKO iPSC cells

WT iPSC cells were seeded in a 6-well plate. On the following day, LNPs containing CRISPR guides were added into the plate. Media was changed daily until cells were split to form single clones. Clones were screened by flow cytometry using HLA-A2 antibody (Invitrogen, clone BB7.2) and HLA-B7 antibody (Miltenyi Biotec, clone REA176) and the ddPCR assays before selecting TKO iPSC clones.

### Generate iPSCs expressing GFP-Luc

Induced pluripotent stem cells (iPSCs) were passaged in mTeSR medium (STEMCELL Technologies) supplemented with 10 µM Rock inhibitor and seeded at a density of 2.5 × 10⁵ cells per well in a 6-well plate pre-coated with Matrigel. The following day, cells were transduced with a lentiviral vector encoding a Luciferase 2A GFP construct under the control of the EF1-α promoter (Tailored Genes, Toronto, ON, Canada) at a multiplicity of infection (MOI) of 5. The medium was replaced 24 h post-transduction with fresh mTeSR. Transduced cells were maintained and expanded in mTeSR medium under standard culture conditions (37°C, 5% CO₂). Following sorting, cells were seeded in Matrigel-coated vessels and cultured in mTeSR for downstream applications. Transduced iPSCs were sorted to isolate GFP-positive cells using the FACSAria II cell sorter (BD Biosciences). Transduction efficiency was assessed one week after transduction by determining the percentage of GFP-positive cells using flow cytometry on a CytoFLEX system (Beckman Coulter). Luciferase activity was quantified using the Promega Luciferase Assay System (Promega, Cat. 1500) following the manufacturer’s protocol.

### Differentiation of iPSCs to pancreatic progenitors or cardiomyocytes

WT and TKO Induced pluripotent stem cells (iPSCs) were differentiated into Pancreatic Progenitor cells using the STEMdiff™ Pancreatic Progenitor Kit (STEMCELL Technologies, Cat. 5120) according to the manufacturer’s protocol, with modifications to the initial seeding density and Matrigel coating procedure. The differentiation efficiency and identity of Pancreatic Progenitor cells were evaluated by Flow Cytometry using NKX6.1 and PDX1 specific antibodies. WT and TKO iPSCs expressing GFP-Luc were differentiated into atrial cardiomyocyte cells using the STEMdiff™ Atrial Cardiomyocyte Differentiation Kit (Stemcell Technologies, Cat. 100- 0215) according to the manufacturer’s instructions, with modifications to the initial seeding density, incubation time for matrigel coating and number of days required to reach >95% confluency for both cell lines. On day 14 of differentiation, cardiac cells were harvested and assessed for expression of cTNT marker (BD Biosciences, Cat. 564767) using flow cytometry.

### *In vitro* MLR assay

Target cells were thawed and seeded into 96-well plates pre-coated with Matrigel (VWR, Cat. CA89050-192) in the presence of Rock Inhibitor (Stemcell Technologies, Cat. 772304). On the second day of culture, HLA expression on target cells was induced by treating the cells with inflammatory cytokines: interferon-gamma (IFN-γ Peprotech, Cat. 300-02, final concentration 100 ng/mL), interleukin-1 beta (IL-1β Peprotech, Cat. 200-01B, final concentration 5 ng/mL), and tumor necrosis factor-alpha (TNF-α Peprotech, Cat. 300-01A, final concentration 10 ng/mL) for 24 h to maximize HLA expression. PBMCs were thawed one day prior to co-culture and cultured overnight in T-cell growth media supplemented with IL-2, IL-7, and IL-15. The following day, PBMCs were harvested, washed, and resuspended in fresh T-cell growth media without cytokine supplementation. On the day of co-culture, PBMCs were added to the target cells at target-to-effector ratios of 1:1. Co-cultures were maintained overnight at 37°C in a humidified incubator with 5% CO₂. The following day, cells were lysed using Promega Luciferase Lysis Buffer (Promega, Cat. E1500). Residual luciferase activity, indicating the viability of target cells, was measured using a plate reader in conjunction with the Promega Luciferase Assay System (Promega, Cat. E1500), following the manufacturer’s instructions. Molecular Device ®, SpectraMax iD5 instrument was used to read Luciferin signals.

## Data availability

The authors declare that all data supporting the findings of this study are available within the paper and its supplementary Information files. Correspondence and requests for materials should be addressed to Y.Z. or A. P.

## Acknowledgements

We sincerely thank Amandeep Dhillon, Louise Moyle, Nabil Zeidan, Pooja Phadiya, Manoja Eswara, Prerna Agrawal, and Elias Uhlin at CCRM/OmniaBio for their contributions to iPSC related experiments and discussions. Their deep expertise and insights in the iPSC field and their effective communication during our productive collaboration are highly appreciated by the authors. We also sincerely thank Christian Dombrowski, Will Harrington, and Ruan Oliveira for contributions relating to the Nme2Cas9 cytosine base editor platform and James Peter for his contributions to the in vivo studies.

## Author contributions

U.J., P.S., I.M., A.L. I.D, Q. Z., B.Z., and O.Z. performed in vitro experiments, analyzed data, produced figures and co-wrote the manuscript. I.B., N.S., and D.L. performed in vivo studies, analyzed bioluminescence imaging experiments, produced figures and co-wrote the manuscript. B.L., and B. H. performed bioinformatic analysis and co-wrote the manuscript. B.S., A.P., and Y.Z. designed experiments, supervised the project, and co-wrote the manuscript.

---

**Sup Fig. 1.**
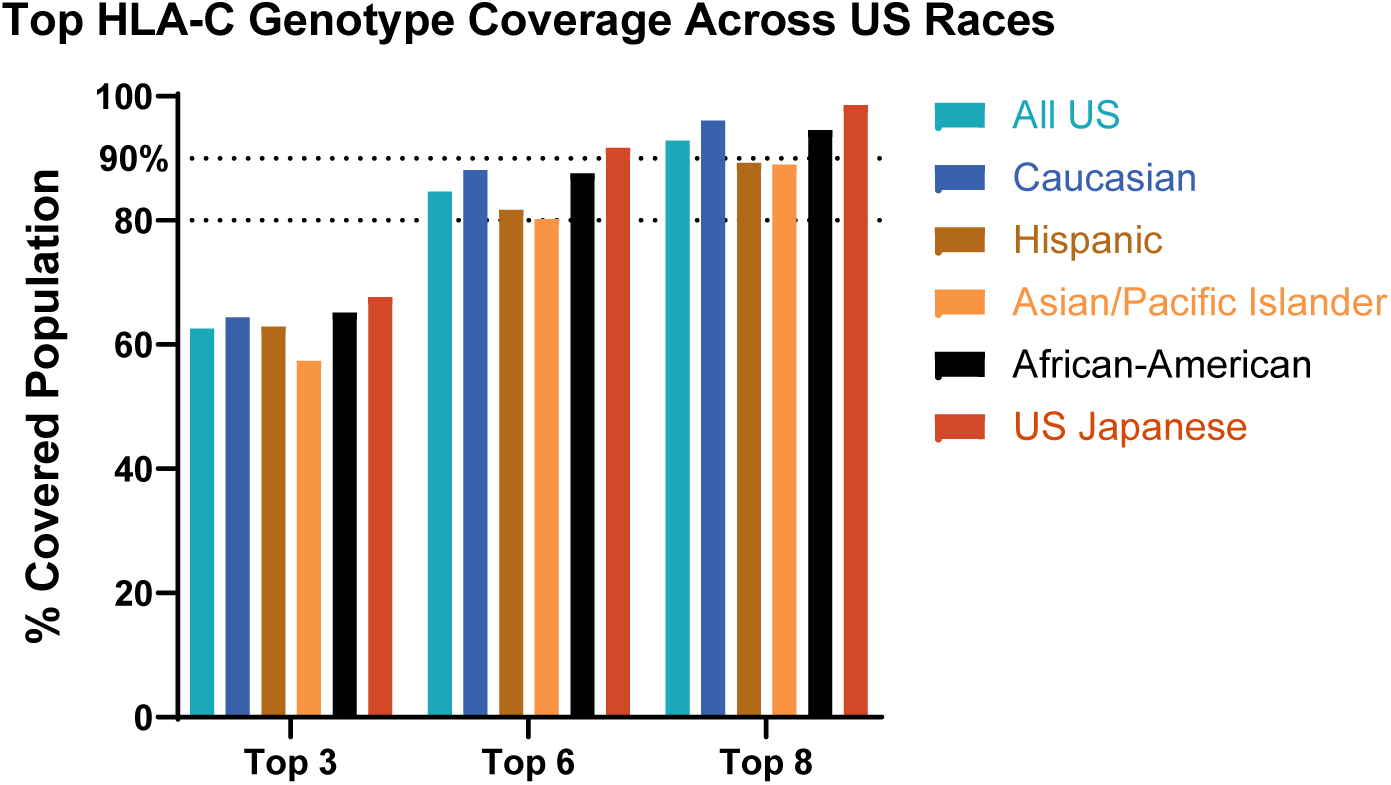
Comparable HLA-C genotype coverage across different US ethnicities. Leveraging a large sample size of over 2 million donors from a US cohort, the subgroup bioinformatic analysis was performed to predict the coverage across different US ethnicities by HLA-C types.

**Sup Fig. 2.**
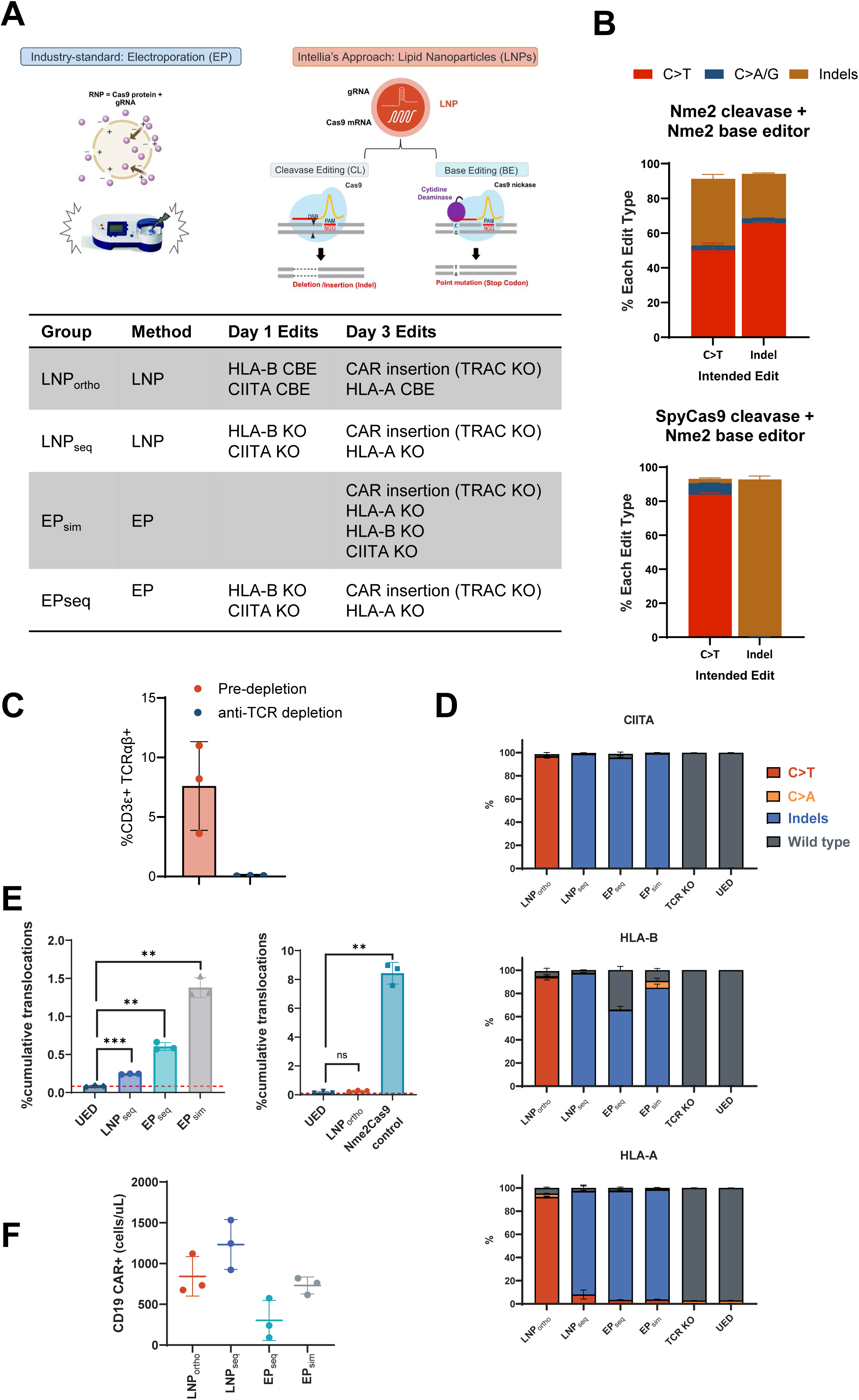
T cell engineering. A. Schematic demonstrating delivery and editing methods commonly used for the ex-vivo T-cell engineering. Table illustrates the editing platform evaluated for each condition described in Fig. 2. B. T cells were multiplex edited with either same species cleavase and base-editor (Nme2Cas9 + Nme2Cas9 CBE), or by using two different Cas9 orthologs (SpyCas9 + Nme2Cas9-CBE) and the type of edit at each site evaluated by amplicon-sequencing. The use of Cas9 orthologs prevents cross-editing at the target sites allowing for controlled combined use of CRISPR/Cas9 base-editing and cleavase editing technologies. C. Edited T cells were depleted for residual endogenous TCR using TCRαβ microbeads achieving >99% TCR removal. D. Confirmation of edit type in each gene target (HLA-A, HLA-B, CIITA) by amplicon-sequencing. E. Quantification of cumulative translocations from ddPCR assays (see Fig. 2). F. Proliferation of Allo-CD19 T cells in *in-vitro* rechallenge assay was measured at round three by quantitative flow cytometry (MACSQuant).

**Sup Fig. 3.**
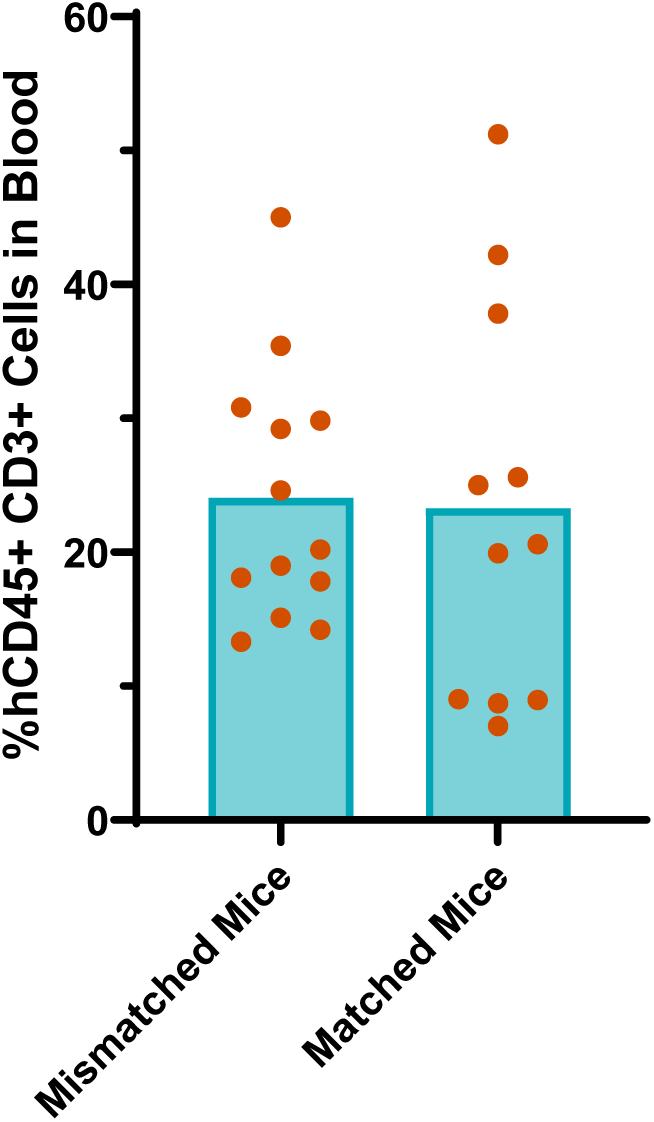
Successful human T cell engraftment in humanized NOG mice. Fifteen weeks after human CD34+ cells were injected into NOG mice, blood samples were collected from the mice and analyzed by flow cytometry using antibodies specific for human CD45 and human CD3. Each dot represents one mouse.

**Sup Fig. 4.**
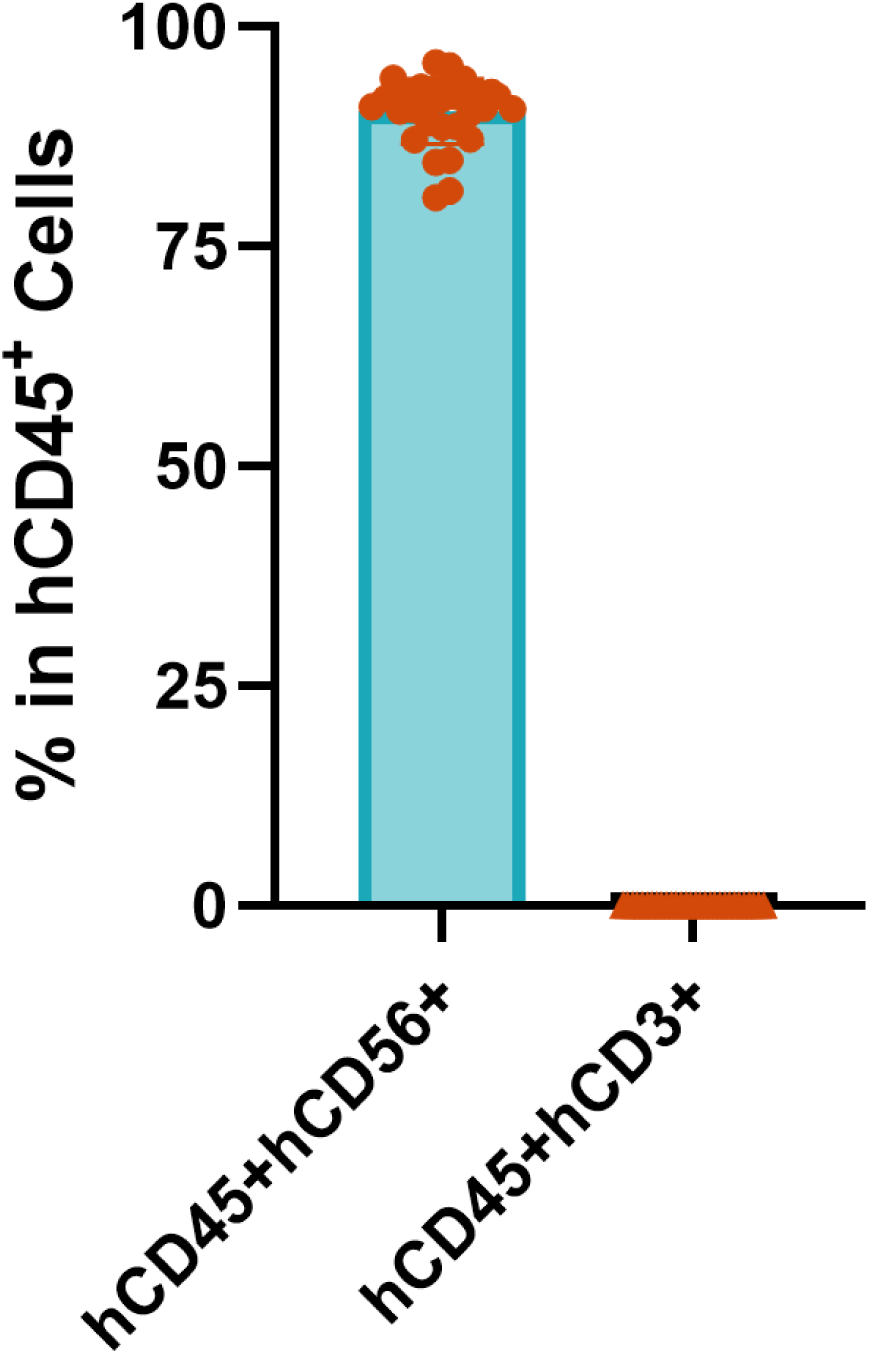
Successful NK engraftment in NOG-IL15 mice. Twenty-nine days after human NK cells were injected into NOG-IL15 mice, blood samples were collected from the mice and analyzed by flow cytometry using antibodies specific for human CD45, human CD56, and human CD3 antibodies. Each dot represents one mouse.

**Sup Fig. 5.**
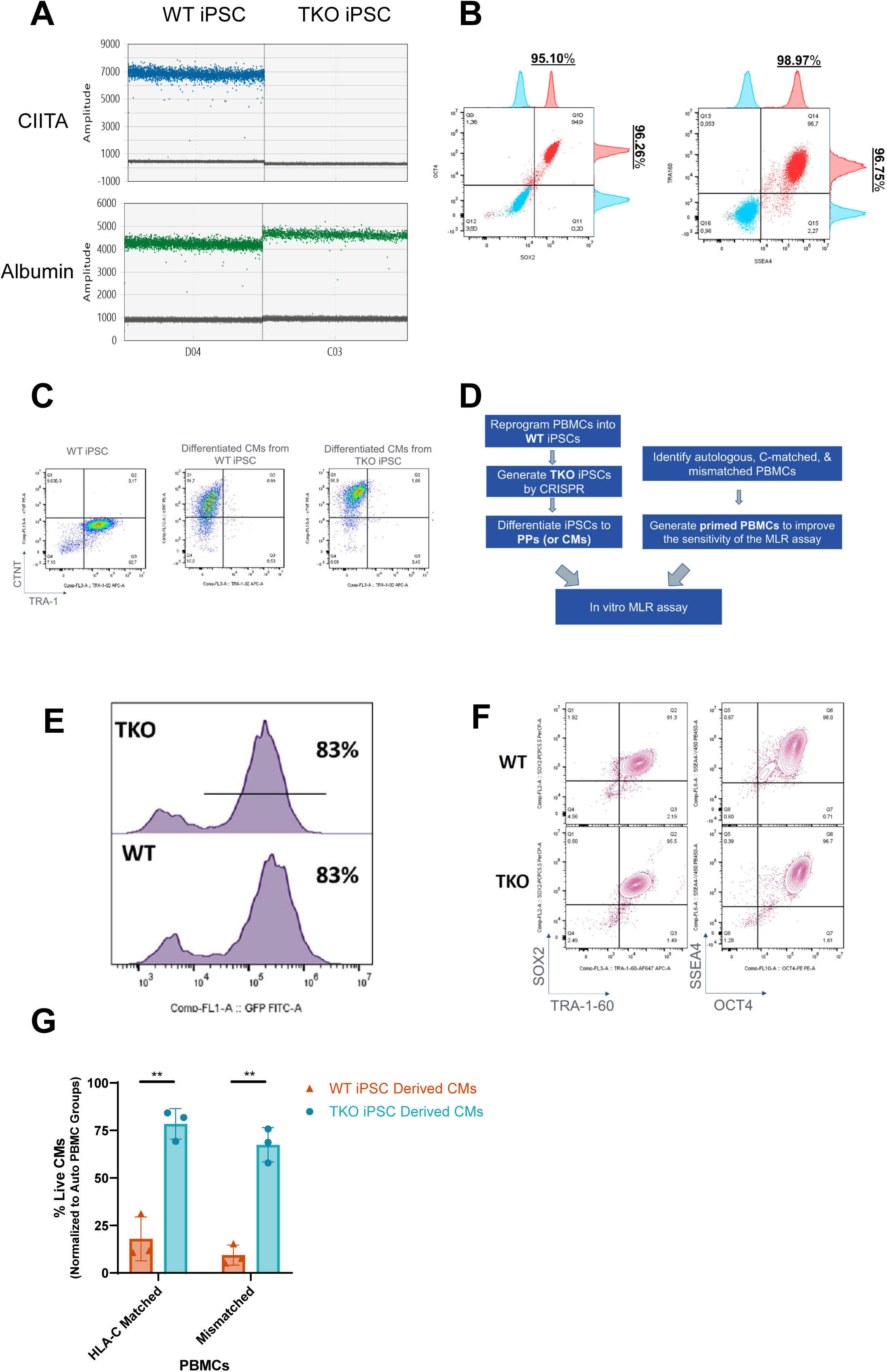
Application of the Allo platform in iPSCs. A. Confirming CIITA KO in TKO iPSCs by the ddPCR assay. B. The expression of pluripotency markers, including SSEA4 and TRA-1-60 (left), and OCT4 and SOX2 (right) in TKO-iPSCs were examined by flow cytometry (red). Samples stained with isotype control antibodies are shown in blue. C. After cardiomyocytes were differentiated from WT or TKO iPSCs, those cells were stained with antibodies to detect a cardiac-specific marker of cTNT and a pluripotent stem cell marker of TRA-1 to confirm cardiomyocyte differentiation. D. Schematic of *in vitro* MLR assay. E. WT or TKO iPSCs were transduced with lentivirus expressing GFP-Luc. Comparable GFP expression level in sorted GFP+ cells were detected by flow analysis. F. Comparable pluripotency marker expression levels were detected in sorted GFP-Luc expressing WT or TKO iPSCs by flow analysis. G. Cardiomyocytes cells differentiated from WT or TKO iPSCs were co-cultured with primed PBMCs from three different hosts, which were autologous host, HLA-C matched host, and mismatched host, at E: T ratio = 1:1. Cardiomyocyte cell lysis data were normalized to data from the autologous host group. Each dot represents one PBMC host. ** *p*<0.01.

